# Destabilization of mRNAs enhances competence to initiate meiosis in mouse spermatogenic cells

**DOI:** 10.1101/2023.09.20.557439

**Authors:** Natalie G. Pfaltzgraff, Bingrun Liu, Dirk G. de Rooij, David C. Page, Maria M. Mikedis

## Abstract

The specialized cell cycle of meiosis transforms diploid germ cells into haploid gametes. In mammals, diploid spermatogenic cells acquire the competence to initiate meiosis in response to retinoic acid. Previous mouse studies revealed that MEIOC interacts with RNA-binding proteins YTHDC2 and RBM46 to repress mitotic genes and promote robust meiotic gene expression in spermatogenic cells that have initiated meiosis. Here, we used the enhanced resolution of scRNA-seq, and bulk RNA-seq of developmentally synchronized spermatogenesis, to define how MEIOC molecularly supports early meiosis in spermatogenic cells. We demonstrate that MEIOC mediates transcriptomic changes before meiotic initiation, earlier than previously appreciated. MEIOC, acting with YTHDC2 and RBM46, destabilizes its mRNA targets, including transcriptional repressors *E2f6* and *Mga*, in mitotic spermatogonia. MEIOC thereby derepresses E2F6- and MGA-repressed genes, including *Meiosin* and other meiosis-associated genes. This confers on spermatogenic cells the molecular competence to, in response to retinoic acid, fully activate transcriptional regulator STRA8-MEIOSIN, required for the meiotic G1/S phase transition and meiotic gene expression. We conclude that in mice, mRNA decay mediated by MEIOC-YTHDC2-RBM46 enhances the competence of spermatogenic cells to initiate meiosis.

**SUMMARY STATEMENT:** RNA-binding complex MEIOC-YTHDC2-RBM46 destabilizes its mRNA targets, including transcriptional repressors. This activity facilitates the retinoic acid-dependent activation of *Meiosin* gene expression and transition into meiosis.

## INTRODUCTION

Sexual reproduction depends on meiosis, a specialized cell cycle that produces haploid gametes via one round of DNA replication followed by two rounds of chromosome segregation. The major chromosomal events of meiosis, including pairing, synapsis, and crossing over of homologous chromosomes, are generally conserved across eukaryotes. By contrast, the mechanisms that govern the transition from mitosis to meiosis are less well conserved (Kimble, 2011). For example, in budding yeast, meiotic initiation occurs when multiple inputs converge to activate a master transcription factor (IME1) that upregulates meiotic gene expression (Kassir et al., 1988; Smith et al., 1990; reviewed in van Werven and Amon, 2011). In *Drosophila melanogaster*, the transition from mitosis to meiosis occurs via translational repression mediated by the RNA helicase Bgcn and its binding partners Bam and Tut (Chen et al., 2014; Chen et al., 2014; Gonczy et al., 1997; Insco et al., 2009; Insco et al., 2012; McKearin and Spradling, 1990).

In mammals, transcriptional activation induced by extrinsic signaling plays a central role in meiotic initiation (i.e., the meiotic G1/S transition). During the transition from mitosis to meiosis (i.e., the “premeiotic” stage during oogenesis and “preleptotene” stage during spermatogenesis), retinoic acid induces meiotic entry by transcriptionally activating *Stra8* and *Meiosin* (Anderson et al., 2008; Baltus et al., 2006; Dokshin et al., 2013; Ishiguro et al., 2020a). *In vitro*, nutrient restriction synergizes with retinoic acid to induce meiotic entry in spermatogenic cells (Zhang et al., 2021). The STRA8-MEIOSIN heterodimer transcriptionally activates expression of G1/S cyclins and meiotic factors that orchestrate the chromosomal events of meiotic prophase I (Anderson et al., 2008; Baltus et al., 2006; Ishiguro et al., 2020a; Kojima et al., 2019; Mark et al., 2008; Soh et al., 2015; Zhang et al., 2023). In mice lacking *Stra8* or *Meiosin* on a C57BL/6 genetic background, premeiotic oogonia and preleptotene spermatocytes halt their development and fail to progress to meiotic prophase I (Anderson et al., 2008; Baltus et al., 2006; Dokshin et al., 2013; Ishiguro et al., 2020a). Indeed, *Stra8*-null premeiotic oogonia and preleptotene spermatocytes, as well as *Meiosin*-null premeiotic oogonia, fail to undergo meiotic DNA replication (Anderson et al., 2008; Baltus et al., 2006; Dokshin et al., 2013; Ishiguro et al., 2020a). (*Meiosin*-null preleptotene spermatocytes initiate an S phase, but it remains unclear whether it is mitotic or meiotic in nature (Ishiguro et al., 2020a).)

In the mammalian testis, competence to initiate meiosis in response to retinoic acid is acquired during spermatogenesis. In mitotic spermatogonia, DMRT1 postpones the acquisition of competence, thereby preventing precocious meiotic entry, by repressing retinoic acid-dependent transcription as well as *Stra8* and likely *Meiosin* gene expression (Matson et al. 2010; Ishiguro et al. 2020). As a result, undifferentiated spermatogonia exposed to endogenous retinoic acid express *Stra8* at low levels, and do not express *Meiosin*, and consequently become mitotic differentiating spermatogonia (Ishiguro et al., 2020a; Matson et al., 2010). Differentiating spermatogonia prematurely exposed to exogenous retinoic acid may express STRA8 protein, but they do not initiate meiosis (Endo et al., 2015; Johnson et al., 2023). At the mitosis-to-meiosis transition, the SCF E3 ubiquitin ligase complex degrades DMRT1 (Matson et al., 2010; Nakagawa et al., 2017), thereby conferring upon spermatogenic cells the competence to express *Stra8* and *Meiosin* and initiate meiosis in response to retinoic acid. To date, the SCF complex is the only known positive regulator of spermatogenic cells’ competence to initiate meiosis.

Post-transcriptional regulation of mRNA also plays a role in governing the mitosis-to-meiosis transition in mammals. After meiotic initiation, MEIOC acts together with RNA helicase YTHDC2 and RNA-binding protein RBM46 to regulate progression through meiotic S phase into meiotic prophase I (Abby et al., 2016; Bailey et al., 2017; Hsu et al., 2017; Jain et al., 2018; Qian et al., 2022; Soh et al., 2017; Wojtas et al., 2017). MEIOC is required to increase meiotic gene expression and also to repress the mitotic cell cycle program, thereby inhibiting a premature and aberrant metaphase several days before wild-type meiotic metaphase I. Consistent with MEIOC, YTHDC2, and RBM46’s homology to *Drosophila* proteins Bam, Bgcn, and Tut, respectively, the MEIOC-YTHDC2-RBM46 complex binds to mRNA (Abby et al., 2016; Bailey et al., 2017; Hsu et al., 2017; Jain et al., 2018; Li et al., 2022; Peart et al., 2022; Qian et al., 2022; Saito et al., 2022; Soh et al., 2017). YTHDC2 interacts with exoribonuclease XRN1, while RBM46 recruits nonsense mediated decay protein UPF1 and subunits of the cytoplasmic deadenylase CCR4-NOT complex, which suggests that YTHDC2 and RBM46’s mRNA targets are degraded (Kretschmer et al., 2018; Li et al., 2022; Qian et al., 2022; Wojtas et al., 2017). The mechanism by which the MEIOC-YTHDC2-RBM46 complex post-transcriptionally regulates its mRNA targets after meiotic initiation remains poorly defined.

Here we molecularly dissect MEIOC’s activity in mouse spermatogenic cells during the mitosis-to-meiosis transition via two parallel approaches: (i) single-cell RNA sequencing (scRNA-seq) analysis of postnatal testes and (ii) bulk RNA-seq analysis of developmentally synchronized testes with histologically verified staging. Both approaches reveal that *Meioc*-null germ cells developmentally diverge from wild-type spermatogenic cells at the meiotic G1/S transition, earlier than previously appreciated. Furthermore, we find that MEIOC’s activity leads to an increase in meiosis-associated gene expression in late mitotic spermatogonia, before meiotic initiation. We discover that MEIOC destabilizes mRNAs encoding transcriptional repressors of meiotic gene expression. This inhibition of repressors facilitates activation of the master meiotic transcriptional regulator STRA8-MEIOSIN in response to retinoic acid. Therefore, a post-transcriptional repressor of mRNA, acting in parallel to the SCF complex’s degradation of DMRT1, enhances spermatogenic cells’ competence to activate the meiotic transcriptional regulator and initiate meiosis.

## RESULTS

### MEIOC drives a transcriptomic shift at the meiotic G1/S transition

To characterize the molecular consequences of MEIOC activity during spermatogenesis, we performed 10x Genomics Chromium-based scRNA-seq on *Meioc*-null and wild-type P15 testes and identified germ cell clusters by cell type-enriched marker expression and transcriptome-based cell cycle analysis (Fig. S1-3). These germ cell clusters, identified in both wild-type and *Meioc*-null testes, consisted of four clusters in mitosis (undifferentiated spermatogonia [Undiff], differentiating types A_1_-A_4_ spermatogonia [A1-4], differentiating Intermediate and type B spermatogonia [In/B], and differentiating type B spermatogonia in G2/M phase [B G2M]); three clusters spanning the mitosis-to-meiosis transition (preleptotene spermatocytes in G1, early S, and late S phase [pL G1, pL eS, and pL lS, respectively]); and three clusters in meiotic prophase I (leptotene spermatocytes [L], zygotene spermatocytes [Z], and pachytene spermatocytes [P]). While prior histological characterization found that *Meioc*-null spermatogenic cells to not progress beyond early zygotene (Abby et al., 2016; Soh et al., 2017), some *Meioc*-null cells were designated as pachytene spermatocytes in the scRNA-seq clustering. This indicates transcriptomic progression in *Meioc*-null spermatocytes is not reflected at the histological level. We also identified a cluster composed of *Meioc*-null spermatocytes, which we designated as “Mutant-only” (Mut). As this cluster was primarily comprised of cells in G2/M phase (Fig. S1C), it likely includes the aberrant metaphase spermatogenic cells found in P15 *Meioc*-null testes, days before meiotic metaphase I occurs in the wild-type testis (Abby et al., 2016; Soh et al., 2017).

We then applied pseudotime analysis to computationally reconstruct the spermatogenic cells’ developmental trajectory from undifferentiated spermatogonia to pachytene spermatocytes (Fig. 1A). *Meioc*-null germ cells followed the same trajectory as wild-type cells during the mitotic stages of spermatogenesis but diverged in the G1 phase of the preleptotene stage, just before meiotic S phase begins. As spermatogenic cells are classically staged via histology, we used histologically-staged samples to independently verify these pseudotime results. We developmentally synchronized spermatogenesis to obtain *Meioc*-null and wild-type testes enriched for preleptotene spermatocytes (Fig. S4A). After histologically verifying staging (Fig. S4B), preleptotene-enriched testes were analyzed via bulk RNA-seq for differential expression, and we identified over 2,000 transcripts as differentially abundant (Fig. S4C). We then asked whether MEIOC impacts the G1/S transition in the preleptotene-enriched testes. We compared percentile ranks for fold changes (WT/*Meioc* KO) for genes enriched at specific cell cycle phases (see Materials and Methods). As a positive control for the disrupted G1/S transition, we re-analyzed a bulk RNA-seq dataset of preleptotene spermatocytes sorted from wild-type and *Stra8*-null testes on a C57BL/6 background (Kojima et al., 2019). This analysis revealed that in preleptotene spermatocytes, MEIOC increased transcript abundance for genes associated with G1/S, S, and G2, similar to STRA8 (Fig. S4D). In total, both scRNA-seq and bulk RNA-seq datasets demonstrate that MEIOC drives a major transcriptomic shift in spermatogenic cells at the meiotic G1/S transition. This is earlier than previously appreciated, as prior histological studies reported that *Meioc*-null spermatogenic cells diverge from wild-type controls after entering meiotic S phase (Abby et al., 2016; Soh et al., 2017).

**Figure 1:**
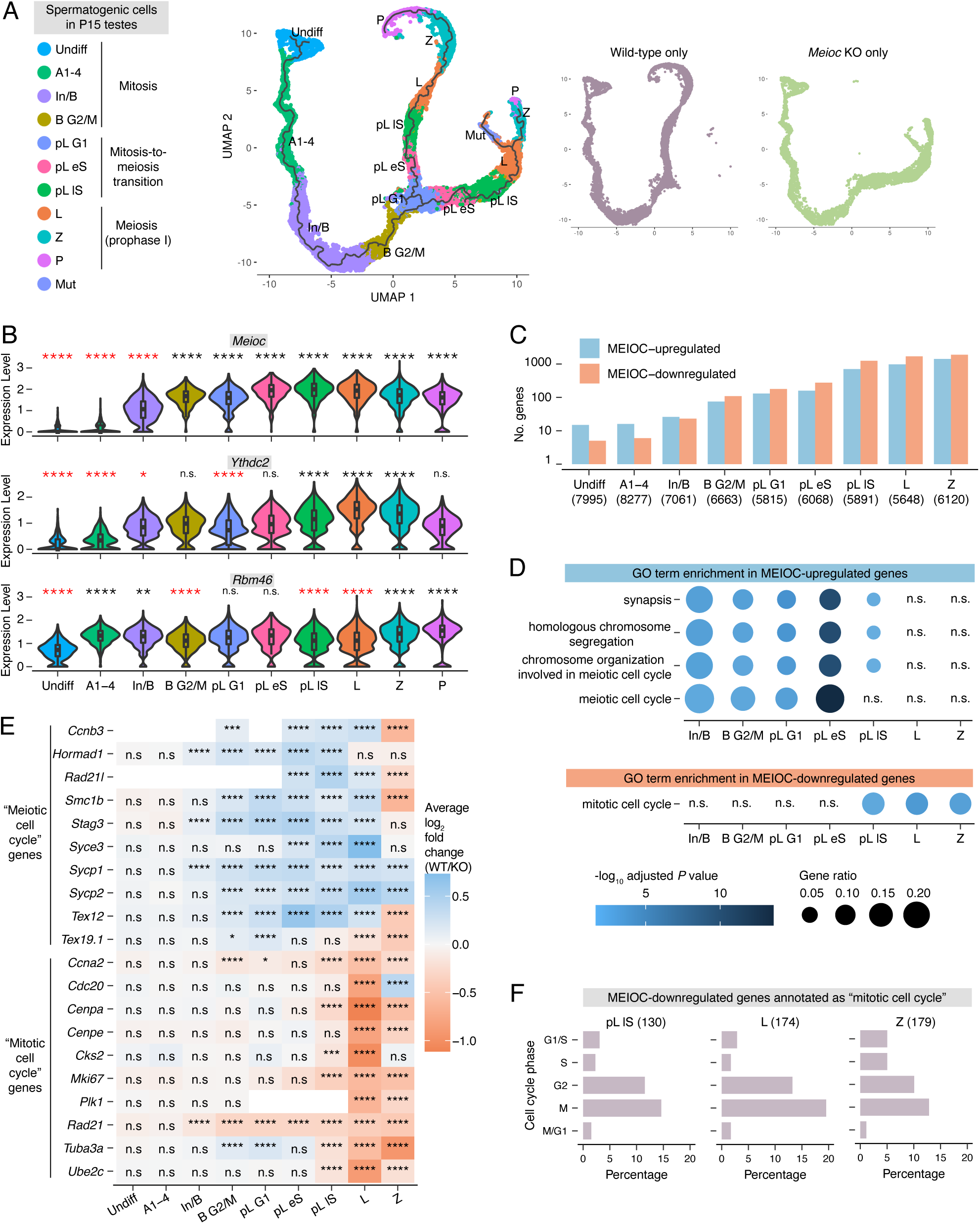
*Meioc*-null germ cells transcriptomically diverge from wild-type germ cells during the G1/S phase transition. **A:** UMAP visualization with pseudotime trajectory of wild-type and *Meioc* KO germ cells from P15 testes. Large plot on left displays both wild-type and *Meioc* KO cells. Smaller plots on right display cells of a single genotype. **B:** Expression levels of *Meioc*, *Ythdc2*, and *Rbm46* in germ cell clusters from wild-type testes. Black and red asterisks designate enrichment or depletion, respectively, relative to all other germ cells. Clusters marked as “not done” (n.d.) did not meet expression thresholds set for statistical testing. **C:** Number of genes identified as MEIOC-upregulated (log_2_ fold change (WT/KO)>0.1, adj. *P*<0.05) and-downregulated (log_2_ fold change (WT/KO)<-0.1, adj. *P*<0.05) within each germ cell cluster. **D:** Gene Ontology (GO) term enrichment analysis for MEIOC-upregulated and -downregulated genes shown. Graph for MEIOC-upregulated genes displays top 4 enriched categories identified in the B G2/M cluster. Graph for MEIOC-downregulated genes displays a selected category of interest. **E:** Heatmap of log_2_ fold changes for selected genes annotated as GO terms “meiotic cell cycle” or “mitotic cell cycle”. **F:** Associated cell cycle phases for MEIOC-downregulated genes that are annotated as GO term “mitotic cell cycle.” Only those genes whose expression is associated with a specific cell cycle phase are graphed. Cluster abbreviations: Undiff, undifferentiated spermatogonia; A1-4, differentiating types A1-A4 spermatogonia; In/B, differentiating Intermediate and type B spermatogonia; B G2M, differentiating type B spermatogonia in G2/M phase; pL G1, preleptotene spermatocytes in G1 phase; pL eS, preleptotene spermatocytes in early S; pL lS, preleptotene spermatocytes in late S phase; L, leptotene spermatocytes; Z, zygotene spermatocytes; P, pachytene spermatocytes; Mut, mutant spermatocytes. *, adj. *P*<0.05; **, adj. *P*<0.01; ***, adj. *P*<0.001; ****, adj. *P*<0.0001; n.s., not significant.

### MEIOC begins to affect the abundance of meiotic, rather than mitotic, transcripts before the meiotic G1/S transition

MEIOC protein and its binding partner YTHDC2 are first detected immunohistochemically at the preleptotene stage, but YTHDC2 protein is also detected via western blotting in mitotic spermatogonia (Abby et al., 2016; Bailey et al., 2017; Hsu et al., 2017; Jain et al., 2018; Soh et al., 2017; Wojtas et al., 2017). Binding partner RBM46 is detected via immunostaining in mitotic spermatogonia through meiotic spermatocytes (Peart et al., 2022; Qian et al., 2022). We found that *Meioc* and *Ythdc2* transcripts became abundant in late mitotic spermatogonia (In/B and B G2M clusters), while *Rbm46* was highly abundant in all germ cell clusters examined (Fig. 1B). These observations raised the possibility that MEIOC, potentially in collaboration with YTHDC2 and RBM46, shapes the germline transcriptome in late mitotic spermatogonia, before the protein can be reliably detected via immunostaining.

To assess how MEIOC impacts the transcriptome during spermatogenesis, we carried out scRNA-seq-based differential expression analysis in germ cell clusters associated with mitotic spermatogonia (Undiff, A1-4, In/B, and B G2M clusters), the mitosis-to-meiosis transition (pL G1, eS, and lS clusters), and meiotic prophase I (L and Z clusters), and identified genes whose transcript abundance is increased (log2 fold change WT/*Meioc* KO > 0.1 and adjusted P < 0.05) or decreased (log2 fold change WT/*Meioc* KO < -0.1 and adjusted P < 0.05) (Fig. 1C) by MEIOC. The number of differentially abundant transcripts increased in the In/B and B G2/M clusters, mirroring *Meioc* and *Ythdc2* expression patterns, and continued to increase thereafter. We conclude that MEIOC begins to affect the germline transcriptome before the meiotic G1/S transition.

Based on functional analysis, transcripts upregulated by MEIOC in the In/B through pL lS clusters were enriched for Gene Ontology (GO) annotations related to meiosis (Fig. 1D). Several transcripts contributing to this enrichment became more abundant in mitotic spermatogonia and remained more abundant through early meiotic spermatocytes (Fig. 1E). However, in the L and Z clusters, these changes in meiosis-associated transcripts no longer represented an enrichment in meiosis-associated GO annotations (Fig. 1D). Bulk RNA-seq analysis of preleptotene-enriched testes also revealed enrichment of meiosis-associated factors among MEIOC-upregulated transcripts (Fig. S4C, E). Therefore, MEIOC activity leads to increased meiotic transcript abundance in late mitotic spermatogonia, earlier than previously appreciated. Misregulation of these transcripts provides a molecular rationale for *Meioc*-null spermatogenic cells’ delayed progression from preleptotene to leptotene to zygotene stages (Abby et al., 2016; Soh et al., 2017).

Given that MEIOC lowers mitotic transcript abundance in fetal oogonia (Soh et al., 2017), we asked whether similar transcriptomic changes were induced by MEIOC in mitotic spermatogonia. MEIOC-downregulated transcripts in the pL lS to Z clusters, but not in developmentally earlier clusters, were enriched for the GO annotation “mitotic cell cycle” (Fig. 1D,E). This included mitotic G1/S and G2/M cyclin *Ccna2* (Fig. 1E), whose downregulation was first evident in B G2/M and pL G1 clusters, before the broader enrichment of mitotic cell cycle-associated transcripts was detected. Consistent with MEIOC’s downregulation of *Ccna2* transcript abundance, *Meioc*-null spermatocytes exhibit prolonged CCNA2 protein expression at leptotene and zygotene stages, when wild-type spermatocytes no longer express this protein (Abby et al., 2016; Soh et al., 2017). Based on a curated list of genes whose expression is linked to specific cell cycle phases (Hsiao et al., 2020), MEIOC-downregulated genes in pL lS to Z clusters were primarily associated with G2 or M phases (Fig. 1F). We conclude that MEIOC’s downregulation of mitotic cell cycle transcripts occurs primarily after the meiotic G1/S transition. The loss of this regulation likely contributes to *Meioc*-null spermatogenic cells’ delayed progression from preleptotene to leptotene to zygotene stages; and their premature progression into an aberrant metaphase state during meiotic prophase I (Abby et al., 2016; Soh et al., 2017).

In sum, MEIOC elevates the abundance of meiosis-associated transcripts beginning in late transit-amplifying spermatogonia through the mitosis-to-meiosis transition and early meiotic prophase I. Then, late in the mitosis-to-meiosis transition into meiotic prophase I, MEIOC broadly lowers mitotic transcript abundance.

### MEIOC destabilizes transcripts that it targets

In early spermatocytes, MEIOC localizes to the cytoplasm and interacts with RNA-binding proteins YTHDC2 and RBM46, which recruit other proteins that degrade mRNA (Kretschmer et al., 2018; Li et al., 2022; Qian et al., 2022; Wojtas et al., 2017). Therefore, we hypothesized that MEIOC degrades its mRNA targets. We first defined transcripts that associate with MEIOC by reanalyzing previously published MEIOC RIP-seq data from P15 testes (Soh et al., 2017), identifying 1,991 MEIOC-bound mRNA (Table S6). We assessed the molecular impact of MEIOC’s interaction with these transcripts via two approaches. First, we examined these targets’ representation among transcripts exhibiting statistically significant abundance changes that were MEIOC-dependent (i.e., MEIOC-upregulated or -downregulated). We found that MEIOC targets were enriched among transcripts whose abundance decreased, but not increased, in response to MEIOC, in the B G2/M through Z clusters (Fig. 2A). Second, we examined all MEIOC targets, irrespective of whether they met a statistical cut-off in the differential expression analysis. MEIOC targets exhibited slightly lower fold changes (WT/KO) than nontargets in the A1-4 through Z clusters (Fig. 2B). An analysis of estimated changes in transcript stability from bulk RNA-seq data showed that MEIOC targets had reduced transcript stability relative to nontargets (Fig. S6A). Therefore, MEIOC lowers the abundance of transcripts that it targets, presumably by promoting their degradation, beginning in mitotic spermatogonia.

**Figure 2:**
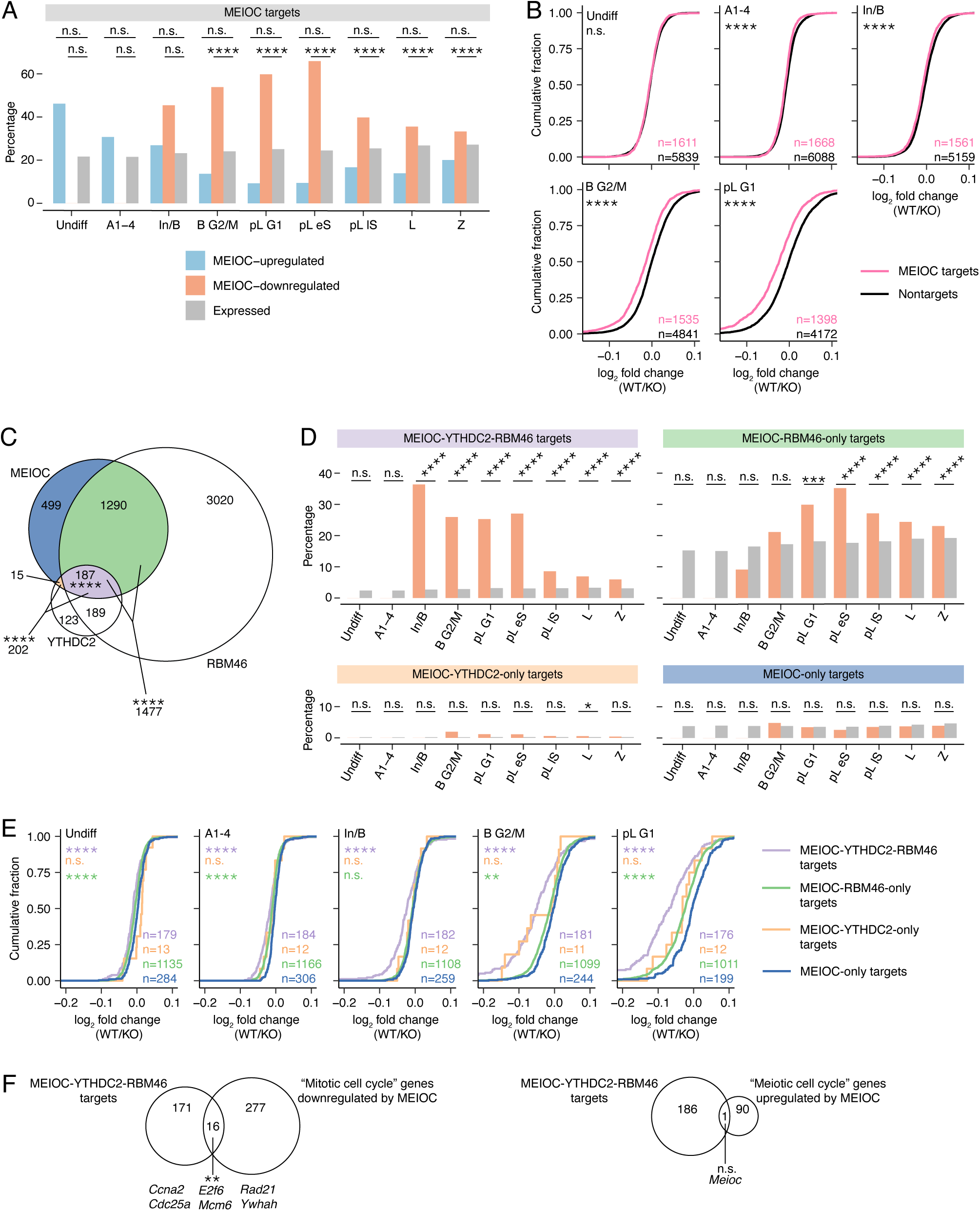
MEIOC-YTHDC2-RBM46 destabilize their mRNA targets. **A:** Percentage of MEIOC targets within MEIOC-upregulated, MEIOC-downregulated, and expressed genes, defined by scRNA-seq. MEIOC targets were identified from reanalysis of a published RIP-seq dataset (Soh et al., 2017). **B:** Cumulative fraction for log_2_ fold change (WT / *Meioc* KO), defined by scRNA-seq, with genes binned as MEIOC targets or nontargets. **C:** Overlap of mRNAs identified by MEIOC RIP-seq, and YTHDC2 CLIP-seq, and RBM46 eCLIP-seq datasets. MEIOC RIP-seq data were reanalyzed from Soh et al., 2017. YTHDC2 CLIP-seq analysis was published in Saito et al., 2022. RBM46 eCLIP-seq data were reanalyzed from Qian et al., 2022. Asterisks denote statistical enrichment. **D:** Percentage of MEIOC-YTHDC2-RBM46 targets, MEIOC-RBM46-only targets, MEIOC YTHDC2-only targets, and MEIOC-only targets within MEIOC-downregulated and expressed genes. **E:** Cumulative fraction for log_2_ fold change (WT / *Meioc* KO), defined by scRNA-seq, with genes binned as MEIOC-YTHDC2-RBM46 targets, MEIOC-RBM46-only targets, MEIOC-YTHDC2-only targets, and MEIOC-only targets. Asterisks represent comparison of color-matched target set to MEIOC-only targets. **F:** MEIOC-YTHDC2-RBM46 targets are enriched for GO term “mitotic cell cycle” genes downregulated by MEIOC but not GO term “meiotic cell cycle” genes upregulated by MEIOC. Upregulated and downregulated genes were defined by scRNA-seq (any germ cell cluster). Asterisks represent statistical enrichment; “n.s.” represents “not significant” statistical enrichment or depletion. *, adj. *P*<0.05; **, adj. *P*<0.01; ***, adj. *P*<0.0001; ****, adj. *P*<0.0001; n.s., not significant.

Given MEIOC’s interaction with YTHDC2 and RBM46, we hypothesized that many MEIOC targets are also bound by YTHDC2 and RBM46. Based on a published YTHDC2 CLIP analysis from testes (Saito et al., 2022) and our reanalysis of RBM46 CLIP data from testes (Qian et al., 2022), MEIOC shared 202 mRNA targets with YTHDC2 and 1,477 targets with RBM46, both of which are statistically significant overlaps (Fig. 2C). The three proteins had 187 mRNA targets in common. Only 15 mRNAs were bound by MEIOC and YTHDC2 but not RBM46, and these likely represent technical differences between datasets rather than a biologically meaningful group of transcripts. MEIOC-YTHDC2-RBM46 targets were enriched for GO annotations associated with RNA stability, while MEIOC-RBM46-only targets did not exhibit statistically significant enrichment (Table S6). We conclude that MEIOC shares many, but not all, of its mRNA targets with YTHDC2 and RBM46.

As YTHDC2 and RBM46 interact with proteins that degrade mRNA, we hypothesized that MEIOC has a greater impact on transcript stability when acting in partnership with YTHDC2 and RBM46 than when acting alone. To test this, we binned MEIOC’s targets as follows: MEIOC-YTHDC2-RBM46 targets, MEIOC-RBM46-only targets, MEIOC-YTHDC2-only, and MEIOC-only targets. We compared how these groups of target genes respond to MEIOC in our scRNA-seq data. First, we examined whether each group was overrepresented among transcripts whose abundance decreased in response to MEIOC. MEIOC-YTHDC2- RBM46 targets exhibited such enrichment beginning in mitotic spermatogonia (Fig. 2D). By contrast, MEIOC-YTHDC2-only targets were enriched in one meiotic cluster and MEIOC-only targets showed no such enrichment (Fig. 2D). MEIOC-RBM46-only targets were enriched beginning at the mitosis-to-meiosis transition (Fig. 2D), suggesting that MEIOC-RBM46 may impact the transcriptome independent of YTHDC2, but the biological significance of this regulation remains unclear. Second, we examined the effect of YTHDC2 and RBM46 on the stability of MEIOC targets, irrespective of statistical cutoff. Relative to MEIOC-only targets, MEIOC-YTHDC2-RBM46 targets exhibited the largest decrease in transcript abundance in all scRNA-seq germ cell clusters (Fig. 2E), as well as in transcript stability in the bulk RNA-seq analysis of preleptotene-enriched testes (Fig. S5B, S6B). Taken together, these data demonstrate that MEIOC’s destabilization of its target mRNAs occurs via its interaction with both YTHDC2 and RBM46.

Given that MEIOC-downregulated transcripts are enriched for mitotic cell cycle factors, we hypothesized that MEIOC-YTHDC2-RBM46 directly targets and destabilizes these factors. By comparing MEIOC-downregulated transcripts bearing the GO annotation “mitotic cell cycle” with MEIOC-YTHDC2-RBM46 targets, we discovered a small but statistically significant overlap that included factors that impact cell cycle progression, such as *Ccna2* and *E2f6* (Fig. 2F). Some targets were downregulated by MEIOC in mitotic spermatogonia and early in the mitosis-to-meiosis transition, before mitotic cell cycle enrichment was evident among the MEIOC-downregulated genes (Fig. 1C,D). Therefore, MEIOC’s destabilization of mRNAs before and during the mitosis-to-meiosis transition may impact spermatogenic cells’ ability to establish a meiosis-specific cell cycle program in meiotic prophase I.

We also considered the possibility that some MEIOC-YTHDC2-RBM46 targets are stabilized, rather than destabilized, by the complex. In particular, given that MEIOC-upregulated transcripts are enriched for the “meiotic cell cycle” annotation (Fig. 1C,D), we asked whether these transcripts were MEIOC-YTHDC2-RBM46 targets. MEIOC-upregulated “meiotic cell cycle” transcripts and MEIOC-YTHDC2-RBM46 targets shared one transcript, *Meioc* (Fig. 2F), with this overlap representing neither statistical depletion nor enrichment. We conclude that MEIOC-YTHDC2-RBM46 does not directly regulate the stability of meiosis-associated transcripts.

In total, MEIOC collaborates with YTHDC2 and RBM46 to promote the decay of the transcripts that it targets, beginning in mitotic spermatogonia.

### MEIOC’s destabilization of *E2f6* and *Mga* transcripts elevates expression of E2F6- and MGA-repressed genes

Most MEIOC-regulated transcripts, as defined by RNA-seq analysis of wild-type versus *Meioc*-null samples, do not directly interact with MEIOC-YTHDC2-RBM46 (Fig. 3A, S6C). We examined whether these changes in transcript abundance reflected altered transcriptional rates or altered transcript stabilities. Applying REMBRANDTS to our bulk RNA-seq data, we estimated changes in mRNA abundance from changes in exonic reads, changes in pre-mRNA abundance (i.e., transcriptional rate) from changes in intronic reads, and changes in mRNA stability from the difference between the changes in exonic and intronic reads (Alkallas et al., 2017; Gaidatzis et al., 2015). MEIOC-regulated transcripts exhibited large changes in transcriptional rates and smaller changes in transcript stabilities (Fig. 3B). We conclude that MEIOC indirectly impacts transcription in ways that alter the abundance of transcripts that it does not bind.

**Figure 3:**
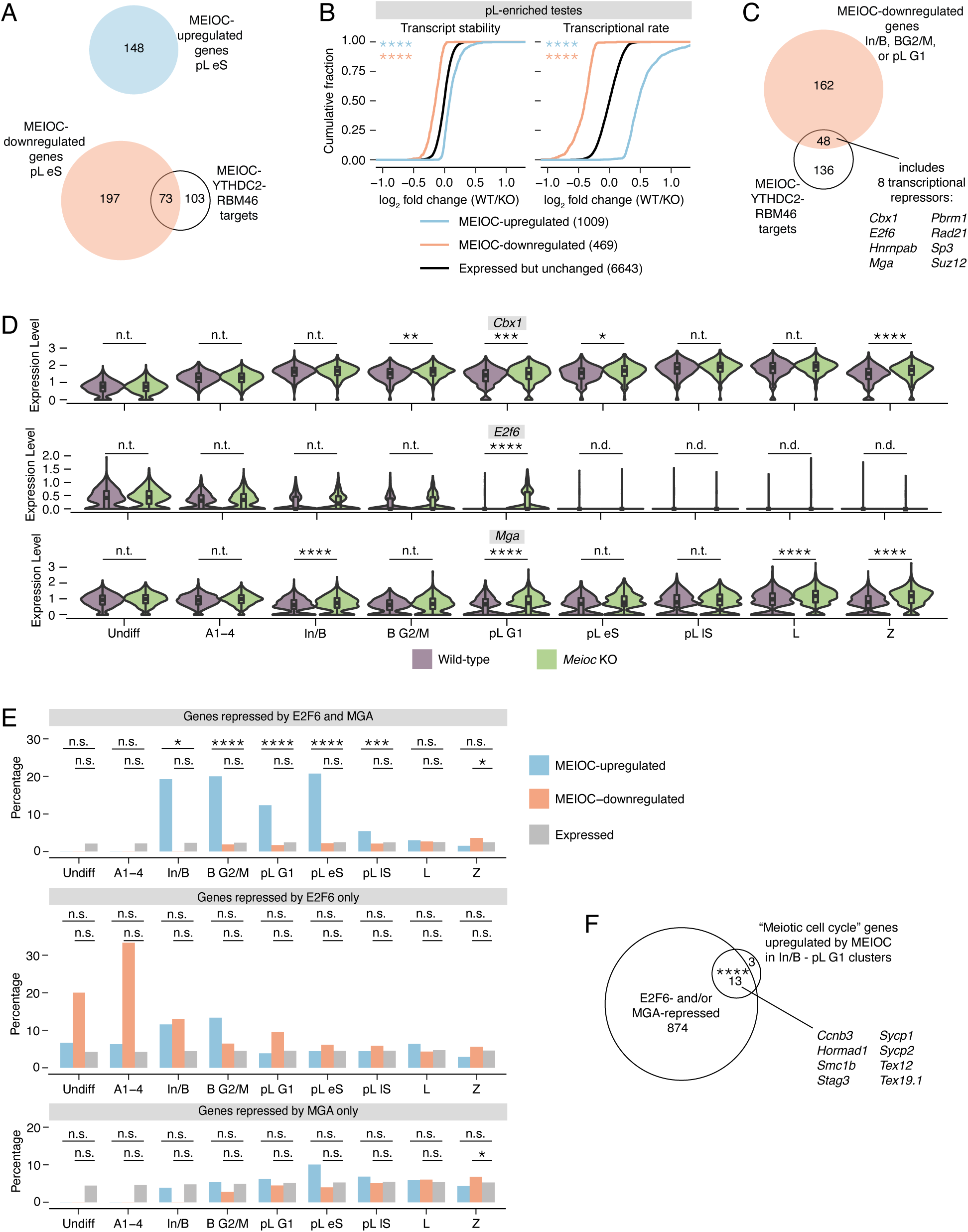
MEIOC-YTHDC2-RBM46’s repression of *E2f6* and *Mga* mRNA relieves E2F6- and MGA-mediated transcriptional repression. **A:** Venn diagram comparing MEIOC-upregulated and -downregulated genes in pL eS cluster to MEIOC-YTHDC2-RBM46 targets. The majority of genes whose transcript abundance changes in response to MEIOC are not directly targeted by MEIOC-YTHDC2-RBM46. **B:** Cumulative distribution of log_2_ fold change (WT/ *Meioc* KO) for transcript stability (left) and transcriptional rate (right) for MEIOC-upregulated and MEIOC-downregulated genes compared to genes that expressed but do not change in response to MEIOC. Transcript stabilities and transcriptional rates were estimated using bulk RNA-seq data from preleptotene-enriched testes. Asterisks represent comparison of color-matched gene set to expressed but not regulated gene set. **C:** Identification of MEIOC-YTHDC2-RBM46 targets that are downregulated by MEIOC in In/B, B G2/M, and/or pL G1 clusters. Of this set, 8 mRNAs encode proteins that have been reported to function as transcriptional repressors. **D:** Expression levels for PRC1.6 subunits *Cbx1*, *E2f6*, and *Mga* in wildtype vs. *Meioc*-null cells in all germ cell clusters identified. *Cbx1*, *E2f6*, and *Mga* are downregulated by MEIOC in the In/B, B G2/M, and/or pL G1 clusters (adj. *P* < 0.05). **E:** Percentage of genes repressed by E2F6 and MGA (top), E2F6 only (center), and MGA only (bottom) among MEIOC-upregulated, MEIOC-downregulated, and expressed genes. E2F6-repressed genes and MGA-repressed genes were identified via reanalysis of published ChIP-seq and RNA-seq datasets from mouse embryonic stem cells (Dahlet et al., 2021; Stielow et al., 2018; Qin et al., 2021). **F:** Overlap between E2F6- and/or MGA-repressed genes and the “meiotic cell cycle” genes upregulated by MEIOC in In/B, B G2/M, and pL G1 clusters. Example genes that fall within this overlap are listed. *, adj. *P*<0.05; **, adj. *P*<0.01; ***, adj. *P*<0.001; ****, adj. *P*<0.0001; n.s., not significant; n.t., not tested (comparison was excluded from statistical testing because log_2_ fold change> -0.1 and <0.1); n.d., not detected (transcript expressed in <25% cells in each population being compared).

We set out to identify MEIOC targets that might drive such transcriptional changes. The vast majority of MEIOC-upregulated transcripts are not directly bound by MEIOC (Fig. 2A, 3A). We hypothesized that MEIOC-YTHDC2-RBM46 destabilizes a mRNA that encodes a transcriptional repressor; destabilization of this mRNA then derepresses (i.e., upregulates) gene expression. For this analysis, we focused on the In/B to pL G1 clusters, before spermatogenic cells have undergone the MEIOC-dependent transcriptomic shift observed in the scRNA-seq pseudotime analysis (Fig. 1A,B). Among 48 MEIOC-YTHDC2-RBM46 targets downregulated by MEIOC in the In/B to pL G1 clusters, we identified eight mRNAs that encode transcriptional repressors: *Cbx1*, *E2f6*, *Hnrnpab*, *Mga*, *Pbrm1*, *Rad21*, *Sp3*, and *Suz12* (Fig. 3C,D). Strikingly, *Cbx1*, *E2f6,* and *Mga* all encode subunits of the noncanonical Polycomb Repressive Complex (PRC) 1.6 (Fig. S7A, S9A), representing a statistically significant enrichment for the complex’s subunits (one-tailed hypergeometric test, *P* = 3.55E-05). *E2f6* and *Mga* are particularly attractive candidates because they encode sequence-specific DNA-binding subunits of PRC1.6 required to repress meiosis-specific genes in somatic and embryonic stem cells (Pohlers et al. 2005; Dahlet et al. 2021; Kehoe et al. 2008; Maeda et al. 2013; Suzuki et al. 2016; Kitamura et al. 2021; Uranishi et al. 2021). By contrast, CBX1 is not strictly required to repress gene expression, as other CBX proteins can compensate for its absence (Ostapcuk et al., 2018).

We hypothesized that MEIOC-YTHDC2-RBM46’s inhibition of *E2f6* and *Mga* impacts gene expression during spermatogenesis. To test this, we reanalyzed ChIP-seq and RNA-seq datasets from mouse embryonic stem cells (Dahlet et al., 2021; Qin et al., 2021; Stielow et al., 2018) to identify genes directly repressed by E2F6 or MGA (see Materials and Methods; Fig. S8A,B). Given that genetic ablation of *Max* (which encodes MGA’s binding partner and a PRC1.6 subunit) induces embryonic stem cells to enter meiosis (Suzuki et al., 2016), these data are relevant to meiotic entry in spermatogenic cells. As E2F6 and MGA cooperate to repress expression of an overlapping set of genes (Dahlet et al., 2021) (Fig. S8C), we classified genes as repressed by both E2F6 and MGA, repressed by E2F6 alone, or repressed by MGA alone. Genes repressed by both E2F6 and MGA, but not those repressed by either factor alone, were overrepresented among MEIOC-upregulated transcripts beginning in the mitotic In/B cluster (Fig. 3E), coinciding with MEIOC’s destabilization of *Mga* mRNA in that cluster (Fig. 3D). This overrepresentation continued through the mitosis-to-meiosis transition (Fig. 3E), including the pL G1 cluster when MEIOC destabilizes *E2f6* and *Mga* mRNA (Fig. 3D). Genes repressed by both E2F6 and MGA were not enriched among MEIOC-downregulated transcripts (Fig. 3E). Bulk RNA-seq analysis of preleptotene-enriched testes largely confirmed these results, while also revealing that MEIOC-upregulated genes were enriched for genes repressed by E2F6 or MGA alone (Fig. S9B), likely because this analysis included lowly expressed genes that were not detected in the sparser scRNA-seq dataset. We conclude that MEIOC’s repression of E2F6 and MGA results in enhanced expression of specific genes in mitotic spermatogonia.

We also examined whether MEIOC’s repression of E2F6 and MGA results in diminished expression of other genes in spermatogenic cells. For this analysis, we used the mouse embryonic stem cell datasets to classify genes as activated by both E2F6 and MGA, by E2F6 alone, or by MGA alone. Genes activated by MGA alone were modestly enriched among MEIOC-downregulated genes within a limited set of clusters (pL lS and L; Fig. S8D). By contrast, genes activated by both E2F6 and MGA or by E2F6 alone were not enriched among MEIOC-upregulated (or downregulated) genes (Fig. S8D). Bulk RNA-seq analysis of preleptotene-enriched testes confirmed these results (Fig. S9B). We conclude that MEIOC’s repression of E2F6 and MGA leads to upregulation of gene expression.

As many MEIOC-upregulated transcripts are meiotic cell cycle factors not bound by MEIOC-YTHDC2-RBM46 (Fig. 1D, 2F), we hypothesized that repression of E2F6 and MGA enhances expression of these meiotic genes. To test this, we asked whether E2F6- and MGA-repressed genes are enriched among meiotic cell cycle transcripts upregulated by MEIOC. We again focused this analysis on the In/B to pL G1 clusters, before spermatogenic cells undergo MEIOC-dependent transcriptomic changes. We found that 13 of 17 MEIOC-upregulated meiotic cell cycle transcripts are targeted for repression by E2F6 and/or MGA (Fig. 3F), and we confirmed these observations in bulk RNA-seq analysis of preleptotene-enriched testes (Fig. S9C). We conclude that MEIOC’s destabilization of *E2f6* and *Mga* mRNAs de-represses meiosis-associated gene expression beginning in mitotic spermatogonia.

### MEIOC indirectly activates transcriptional regulator STRA8-MEIOSIN to enhance spermatogenic cells’ competence to initiate meiosis

In mouse embryonic stem cells, E2F6 and MGA directly repress *Meiosin* (*Gm4969*, *Bhmg1*) (Uranishi et al., 2021) (Fig. S8A,B). Consistent with MEIOC’s destabilization of *E2f6* and *Mga* mRNAs, we found that MEIOC enhances *Meiosin* expression at both the transcript and protein levels during the mitosis-to-meiosis transition (Fig. 4A, S4C, S10A). By meiotic prophase I, *Meioc*-null germ cells exhibited delayed upregulation of *Meiosin* expression (Fig. 4A). We confirmed that *Meiosin* expression is also increased by YTHDC2 and RBM46, based on our examination of published RNA-seq data from postnatal testes (Jain et al., 2018; Peart et al., 2022). MEIOC protein does not bind *Meiosin* mRNA (Table S6). This suggests that MEIOC-YTHDC2-RBM46’s regulation of *E2f6* and *Mga* may be impacting *Meiosin* gene expression.

**Figure 4:**
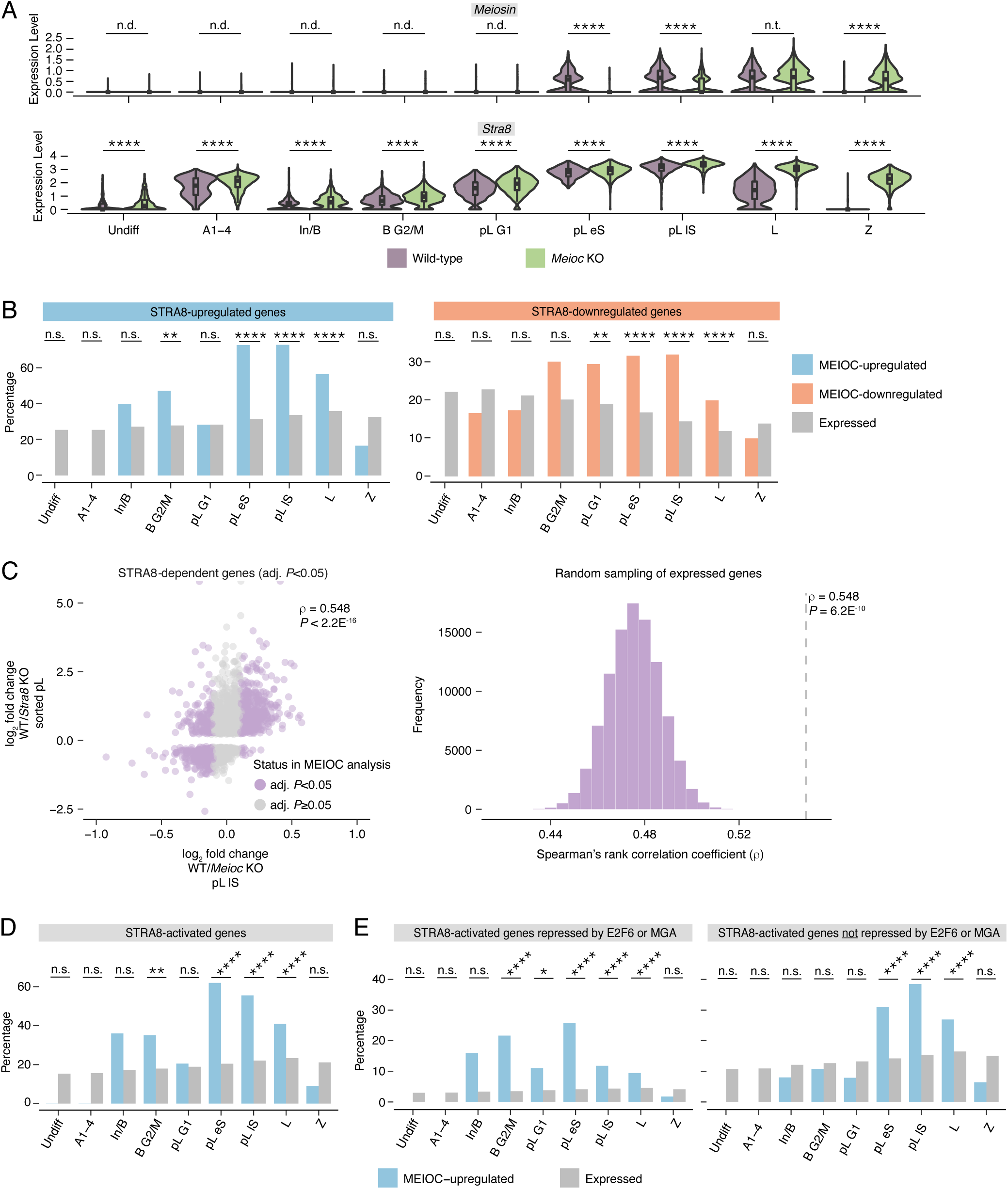
MEIOC’s derepression of *Meiosin* gene expression enhances activation of the STRA8-MEIOSIN transcriptional program. **A:** Expression levels of *Meiosin* and *Stra8* in wild-type and Meioc KO in all germ cell clusters. Clusters marked as “not done” (n.d.) did not meet expression thresholds set for statistical testing. **B:** Percentage of STRA8-upregulated and -downregulated genes in MEIOC-upregulated, - downregulated, and expressed genes from scRNA-seq analysis. STRA8-upregulated and - downregulated genes were identified via reanalysis of bulk RNA-seq data from wild-type and Stra8 KO sorted preleptotene spermatocytes from Kojima et al., 2019. **C:** Left panel, correlation between MEIOC scRNA-seq analysis of the pL lS cluster and STRA8 bulk RNA-seq analysis of sorted preleptotene spermatocytes. Analysis was limited to genes that were statistically dependent on STRA8 (adj. *P*<0.05). *P* value represents the probability that Spearman rho does not equal 0. Right panel, distribution of correlations for gene sets obtained by random sampling of genes expressed in the scRNA-seq pL lS cluster and bulk RNA-seq sorted preleptotene spermatocytes. *P* value represents that probability of obtaining an equal or larger correlation by random sampling. **D:** Percentage of STRA8-activated genes in MEIOC-upregulated and expressed genes from scRNA-seq analysis. STRA8-activated genes were identified as those genes with STRA8-bound promoters (as identified by Kojima et al. (2019) via STRA8-FLAG ChIP-seq in preleptotene-enriched testes) and upregulated by STRA8 (as identified by reanalysis of bulk RNA-seq data from wild-type and Stra8 KO sorted preleptotene spermatocytes from Kojima et al., 2019). **E:** Percentage of STRA8-activated genes repressed by E2F6 or MGA (left), as well as STRA8-activated genes not repressed by E2F6 or MGA (right), in MEIOC-upregulated and expressed genes. *, adj. *P*<0.05; **, adj. *P*<0.01; ***, adj. *P*<0.001; ****, adj. *P*<0.0001; n.s., not significant; n.t., not tested (comparison was excluded from statistical testing because log_2_ fold change> -0.1 and <0.1); n.d., not detected (transcript expressed in <25% cells in each population being compared).

We examined whether MEIOC affects other known regulators of *Meiosin* gene expression. As retinoic acid transcriptionally activates *Meiosin* (Ishiguro et al., 2020b), we considered whether MEIOC enhances retinoic acid-mediated transcription, but we found no evidence for this model. MEIOC did not increase the transcript abundance of any retinoic acid receptor (RAR) or retinoid X receptor (RXR), which mediate transcriptional activation by retinoic acid (reviewed in Endo et al., 2019) (Fig. S10B,C). In addition, MEIOC did not upregulate additional retinoic acid-activated genes *Stra8*, which encodes MEIOSIN’s binding partner, and *Rec8*, a meiotic cohesin (Koubova et al., 2014; Soh et al., 2015; Zhang et al., 2021) (Fig. 4A, S10C,D). We observed that MEIOC downregulated *Stra8* abundance overall in the scRNA-seq data (Fig. 4A), but as *Stra8* mRNA is not directly bound by MEIOC (Table S6), the molecular basis for this regulation remains uncharacterized. These observations indicate that MEIOC does not activate *Meiosin* gene expression by increasing retinoic acid-mediated transcription.

We also considered whether *Dmrt1*, whose encoded protein presumably represses *Meiosin* gene expression (Ishiguro et al., 2020b), was regulated by MEIOC-YTHDC2-RBM46, but again, we found no evidence for this alternative model. *Dmrt1* transcript was not a target of MEIOC, YTHDC2, or RBM46 (Table S6), nor was it differentially expressed in response to MEIOC in the pL G1 or pL eS clusters of the scRNA-seq analysis or in the preleptotene-enriched testes of the bulk RNA-seq analysis (Fig. S11A,B). DMRT1 protein expression was similar in both wild-type and *Meioc*-null testes, with DMRT1 expressed in mitotic spermatogonia but absent from preleptotene spermatocytes (Fig. S11C). In addition, MEIOC did not affect the expression of DMRT1-regulated genes *Tbx1* and *Crabp2* (Matson et al., 2010) (Fig. S11D,E). DMRT1 also directly inhibits *Stra8* expression (Matson et al., 2010), but as noted above, *Stra8* was not activated by MEIOC (Fig. 4A). Therefore, changes in DMRT1 do not account for MEIOC’s upregulation of *Meiosin* gene expression. Taking these observations together, we conclude that MEIOC’s repression of *E2f6* and *Mga* derepresses *Meiosin* gene expression during the mitosis-to-meiosis transition.

STRA8-MEIOSIN functions as an obligate heterodimer that transcriptionally activates gene expression during the mitosis-to-meiosis transition (Ishiguro et al., 2020a; Kojima et al., 2019). As described above, *Stra8* is highly expressed at both the transcript and protein levels in *Meioc*-null spermatogenic cells at the mitosis-to-meiosis transition (Fig. 4A, S4C, Fig. S10A), as previously reported (Abby et al., 2016; Soh et al., 2017). We hypothesized that MEIOC’s upregulation of *Meiosin* gene expression leads to STRA8-MEIOSIN-mediated transcriptional changes. We tested three predictions of this hypothesis.

First, we tested whether, during the mitosis-to-meiosis transition, genes dependent on MEIOC also depend on STRA8. We defined STRA8-dependent (i.e., STRA8-upregulated or - downregulated) genes using bulk RNA-seq data from wild-type and *Stra8*-null preleptotene spermatocytes isolated via synchronization and sorting (Kojima et al., 2019). STRA8-upregulated genes were enriched among MEIOC-upregulated genes during the mitosis-to-meiosis transition (pL eS and lS clusters) and in meiotic prophase I (L cluster; Fig. 4B). Similarly, STRA8-downregulated transcripts were enriched among MEIOC-downregulated transcripts at these same stages (Fig. 4B). Bulk RNA-seq analysis of preleptotene-enriched testes produced similar results (Fig. S12A).

Second, focusing exclusively on STRA8-dependent genes, we looked for correlated changes in transcript abundance in the MEIOC scRNA-seq and STRA8 bulk RNA-seq datasets. STRA8-dependent genes exhibited significantly correlated effects in the MEIOC and STRA8 datasets that peaked at the mitosis-to-meiosis transition (pL eS and lS clusters; Fig. 4C, S13A, S14A). STRA8-dependent genes similarly exhibited robustly correlated effects in the MEIOC bulk RNA-seq dataset from preleptotene-enriched testes and the STRA8 dataset (Fig. S12B). We conclude that MEIOC activity leads to STRA8-MEIOSIN-mediated changes in transcript abundance.

Third, we tested whether MEIOC-dependent genes are directly activated by STRA8-MEIOSIN. We identified STRA8-activated genes as those genes (i) whose expression increased in response to STRA8 and (ii) whose promoters were bound by STRA8 (as defined by ChIP-seq from testes enriched for preleptotene spermatocytes (Kojima et al., 2019)). We found that MEIOC-upregulated genes were enriched for STRA8-activated genes during the mitosis-to-meiosis transition (pL eS and lS clusters; Fig. 4D). However, de novo motif analysis failed to identify enrichment of any motif among these MEIOC-upregulated genes, likely due to the sparsity of scRNA-seq data. The enrichment of STRA8-activated genes among MEIOC-upregulated genes was also visible in the bulk RNA-seq analysis of preleptotene-enriched testes

(Fig. S12C). Furthermore, the top motif identified by de novo motif analysis of MEIOC-upregulated genes from the bulk RNA-seq analysis matched the STRA8-MEIOSIN binding motif (Fig. S12D). The second top motif matched the binding site shared by E2F6 and other E2F proteins (Fig. 12D), as previously reported among STRA8-activated genes (Kojima et al., 2019). In total, ∼60-70% of MEIOC-upregulated genes are directly activated by STRA8-MEIOSIN.

Together, these analyses demonstrate that MEIOC indirectly activates the STRA8-MEIOSIN transcriptional regulator, thereby facilitating a massive shift in the transcriptome at the meiotic G1/S transition (Fig. 1A). With this new molecular insight, we revisited our interpretation of the *Meioc*-null phenotype. Based on prior histological analyses and DNA labeling experiments, *Meioc*-null spermatogenic cells successfully enter the preleptotene stage, express STRA8 protein, and exhibit DNA synthesis. However, they then accumulate at the preleptotene stage and are delayed in entering meiotic prophase I (Abby et al., 2016; Soh et al., 2017). Previously, we interpreted these data as *Meioc*-null spermatogenic cells successfully undergoing the meiotic G1/S phase transition but thereafter exhibiting disrupted progression into meiotic prophase I. Given our present findings that *Meioc*-null germ cells are delayed in upregulating *Meiosin* expression and exhibit a disrupted meiotic G1/S phase transition, we now interpret the *Meioc*-null phenotype as defective meiotic initiation. We conclude that MEIOC enhances spermatogenic cells’ competence to initiate meiosis by activating the STRA8-MEIOSIN transcriptional regulator.

### MEIOC derepresses meiotic gene expression before activating STRA8-MEIOSIN

We observed that many STRA8-activated genes are repressed by E2F6 and/or MGA (Fig. S12E). Consistent with this observation, STRA8-activated genes are enriched for the binding motif of E2F6 and other E2F proteins (Fig. 12D; Kojima et al., 2019). Accordingly, we divided STRA8-activated genes into two groups, based on whether or not they were directly repressed by E2F6 and/or MGA in embryonic stem cells. As MEIOC represses E2F6 and MGA in mitotic spermatogonia and activates STRA8-MEIOSIN during the mitosis-to-meiosis transition, we predicted that genes experimentally defined as E2F6 and/or MGA-repressed and STRA8-activated would be upregulated by MEIOC starting in mitotic spermatogonia, coincident with MEIOC’s destabilization of *E2f6* and *Mga* mRNAs. Conversely, STRA8-activated genes not repressed by E2F6 or MGA should be upregulated by MEIOC later in the mitosis-to-meiosis transition, coincident with MEIOC-mediated upregulation of *Meiosin* gene expression. Consistent with these predictions, STRA8-activated, E2F6-MGA-repressed genes were enriched among MEIOC-upregulated genes in the mitotic B G2/M cluster and during the mitosis-to-meiosis transition (pL G1, eS, and lS clusters; Fig. 4E). Also as predicted, STRA8-activated genes not repressed by E2F6 or MGA exhibited enrichment in the pL eS and lS clusters (Fig. 4E). Both gene sets were also enriched among MEIOC-upregulated genes in the bulk RNA-seq analysis (Fig. S12F). We conclude that, during late transit amplification, MEIOC derepresses a set of meiotic genes targeted by E2F6 and MGA; then during the mitosis-to-meiosis transition, MEIOC elevates expression of these and other meiotic genes by activating STRA8-MEIOSIN.

In total, MEIOC-YTHDC2-RBM46’s destabilization of *E2f6* and *Mga* mRNAs in late mitotic spermatogenic cells derepresses E2F6-MGA-targeted genes, including *Meiosin* and other meiosis-associated genes. In turn, these steps drive activation of the STRA8-MEIOSIN transcriptional regulator by retinoic acid during the mitosis-to-meiosis transition (Fig. 5). As STRA8-MEIOSIN is the molecular vehicle through which retinoic acid induces the meiotic G1/S transition, MEIOC’s regulation of the STRA8-MEIOSIN complex enhances spermatogenic cells’ competence to initiate meiosis.

**Figure 5:**
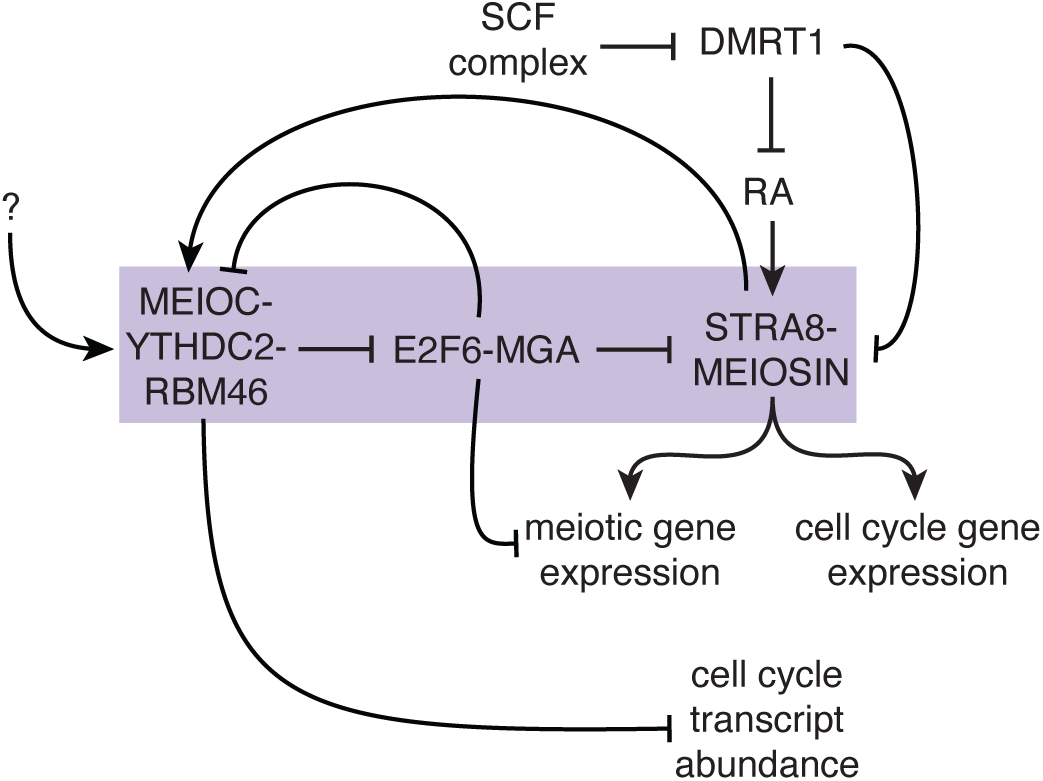
Model for MEIOC-YTHDC2-RBM46 enhancing spermatogenic cells’ competence to transition from mitosis to meiosis in response to retinoic acid. MEIOC-YTHDC2-RBM46 destabilize *E2f6* and *Mga* mRNA and thereby inhibit E2F6 and MGA’s repression of transcription at genes involved in meiosis, including *Meiosin* and *Meioc*. In parallel, the SCF complex degrades DMRT1 and consequently inhibits DMRT1’s repression of retinoic acid (RA)-dependent transcription as well as *Stra8* and *Meiosin* gene expression. Retinoic acid activates *Stra8* and *Meiosi*n gene expression. This activates the STRA8-MEIOSIN transcription factor, which drives the transcription of cell cycle genes as well as meiotic genes, many of which were previously repressed by E2F6 and MGA. MEIOC’s repression of cell cycle transcripts also contributes to the establishment of a meiosis-specific cell cycle program in meiotic prophase I. The question mark denotes undefined molecular regulation that activates MEIOC-YTHDC2-RBM46 before retinoic acid activates STRA8-MEIOSIN transcriptional activity and the transition from mitosis to meiosis. Purple box highlights the novel regulation identified in this study that facilitates competence for the mitosis-to-meiosis transition.

## DISCUSSION

Here we demonstrate that *Meioc*-null spermatogenic cells developmentally diverge from their wild-type counterparts during the meiotic G1/S transition, earlier than previously appreciated. MEIOC is required to derepress expression of meiotic genes, including the transcriptional regulator *Meiosin*. In turn, MEIOC elevates expression of genes targeted by STRA8-MEIOSIN, which drives meiotic initiation. Therefore, MEIOC enhances spermatogenic cells’ competence to activate the meiotic transcriptional regulator and initiate meiosis in response to retinoic acid.

We find that MEIOC-YTHDC2-RBM46 destabilizes its mRNA targets, as suggested previously based on analyses of transcript abundance (Hsu et al., 2017; Qian et al., 2022; Saito et al., 2022; Soh et al., 2017). Here we distinguished changes in transcription vs. transcript stability via two approaches. First, scRNA-seq allowed us to identify and analyze spermatogenic cells impacted by MEIOC before the onset of major transcriptional changes. Second, we employed a specialized pipeline to distinguish between transcriptional rate and RNA stability in bulk RNA-seq data of preleptotene-enriched testes (Alkallas et al., 2017). These complementary methods confirmed that MEIOC-YTHDC2-RBM46 reduces the stability of its target transcripts. Transcript stability may be the primary mechanism by which MEIOC-YTHDC2-RBM46 regulates its targets, as a recent study using ribosome profiling found that YTHDC2 does not affect translation in postnatal testes (Saito et al., 2022).

MEIOC-YTHDC2-RBM46 elevates the abundance of meiosis-associated transcripts without binding them. Here we demonstrated that these transcript abundance changes are driven by changes in transcription (Fig. 3B). Further, we provide a two-step mechanism for these indirect effects on transcript abundance. First, MEIOC-YTHDC2-RBM46 binds to and destabilizes *E2f6* and *Mga* mRNAs, which encode transcriptional repressors whose genomic targets are enriched for meiosis-associated genes (Dahlet et al., 2021; Kehoe et al., 2008; Kitamura et al., 2021; Maeda et al., 2013; Pohlers et al., 2005; Suzuki et al., 2016; Uranishi et al., 2021). This regulation impacts gene expression, and potentially differentiation, beginning in late mitotic spermatogonia, when MEIOC begins to destabilize *Mga* (Fig. 3D). Second, this MEIOC-YTHDC2-RBM46-mediated repression of *E2f6* and *Mga* derepresses *Meiosin* gene expression and activates STRA8-MEIOSIN. This transcriptional regulator then activates meiotic gene expression and the meiotic G1/S transition. Thus, MEIOC-YTHDC2-RBM46’s repression of a transcriptional repressor confers developmental competence to initiate meiosis.

MEIOC is also required in meiotic oocytes for progression through early meiotic prophase I (Abby et al., 2016; Soh et al., 2017), and it remains an open question whether MEIOC supports activation of STRA8-MEIOSIN during meiotic initiation in premeiotic oogonia. There are some molecular differences in meiotic initiation between oogonia and spermatogenic cells. At stages when wild-type germ cells are undergoing meiotic DNA replication, *Stra8*-null oogonia fail to initiate any DNA replication, while *Stra8*-null spermatogenic cells initiate a DNA replication that is non-meiotic in nature, as meiotic cohesin REC8 is not loaded onto the chromosomes (Anderson et al., 2008; Baltus et al., 2006; Dokshin et al., 2013). STRA8 has the additional function of sequestering RB1, a key regulator of the G1/S transition, to promote a timely meiotic G1/S transition in oogonia, but this activity does not appear to impact spermatogenesis (Shimada et al., 2023).

In mitotic oogonia, loss of PRC1 activity via genetic ablation of *Ring1* and *Rnf2* causes premature meiosis due to precocious expression of *Stra8* and other meiosis-associated genes (Yokobayashi et al., 2013). In addition, genetic ablation of *Max,* which encodes a subunit of PRC1.6, induces a similar phenotype with precocious expression of *Stra8*, *Meiosin*, and meiosis-associated genes (Suzuki et al., 2024). During spermatogenesis, PRC1.6 may also regulate gene expression and meiotic initiation, but the germline roles for PRC1.6, E2F6, or MGA remain uncharacterized. While *E2f6*-null mice are fertile, spermatogenesis was reportedly disrupted, without detailed characterization (Storre et al., 2002). Genetic ablation of *Mga* causes embryonic lethality, precluding analysis of spermatogenesis (Burn et al., 2018; Washkowitz et al., 2015). Further studies will be required to characterize E2F6 and MGA’s roles in spermatogenesis, including whether loss of E2F6 and MGA rescues the developmental competence of *Meioc*-null spermatogenic cells. Other transcriptional repressors targeted by MEIOC-YTHDC2-RBM46 may also control *Meiosin* gene expression.

As some *Meioc*-null spermatogenic cells upregulate *Meiosin* gene expression and enter meiotic prophase I on a delayed timeline, MEIOC is not strictly required for competence to initiate meiosis. Perhaps residual activity of YTHDC2-RBM46 in the absence of MEIOC can support meiotic initiation in some cells. Consistent with this possibility, YTHDC2 protein is still expressed in *Meioc*-null spermatogenic cells (Abby et al., 2016; Soh et al., 2017), but whether loss of MEIOC affects RBM46 protein expression remains unclear. Alternatively, acquisition of competence mediated by the SCF complex may enable meiotic initiation on this delayed timeline. Regardless, we have found that the *Meioc*-null, *Ythdc2*-null, or *Rbm46*-null phenotype of delayed entry into meiotic prophase I is the result of reduced STRA8-MEIOSIN transcriptional activation and is less severe than the arrest at the G1/S transition, before meiotic prophase I, exhibited by *Stra8*-null or *Meiosin*-null spermatogenic cells on an inbred C57BL/6 background. Intriguingly, *Stra8*-null spermatogenic cells on a mixed genetic background exhibit the less severe phenotype seen with *Meioc*-null, *Ythdc2*-null, or *Rbm46*-null spermatogenic cells. Our analyses of the *Meioc*-null phenotype suggest that, on a mixed genetic background, *Stra8*-null spermatogenic cells transcriptionally activate some meiotic gene expression, perhaps due to MEIOSIN acting as a homodimer, but further studies will be required to test this possibility.

Our model of mammalian meiotic initiation exhibits parallels to the molecular network that governs meiotic initiation in budding yeast. In yeast, inhibition of a transcriptional repressor (Rme1p) via mRNA destabilization activates expression of the key transcription factor (Ime1p) that governs meiotic entry. We conclude that in both unicellular eukaryotes and multicellular organisms, destabilization of a transcriptional repressor at the transcript level controls activation of the meiotic transcriptional program.

We have placed transcriptional activation by STRA8-MEIOSIN and post-transcriptional repression of mRNA by MEIOC-YTHDC2-RBM46 in a positive feedback loop that facilitates the mitosis-to-meiosis transition (Fig. 5). This model generates a new question: how is MEIOC-YTHDC2-RBM46 activated before retinoic acid activates STRA8-MEIOSIN, particularly when E2F6 and MGA repress *Meioc* gene expression (Fig. 5)? One possibility is illustrated by MEIOC’s *Drosophila* homolog Bam, which (translationally) represses its mRNA targets in late mitotic spermatogonia to facilitate the transition from mitosis to meiosis (Insco et al., 2009; Insco et al., 2012). This transition requires that Bam protein accumulates to a critical threshold level (Insco et al., 2009). Whether mammalian MEIOC protein levels must also clear a critical threshold to activate the MEIOC-YTHDC2-RBM46 complex requires further investigation.

In conclusion, by destabilizing its mRNA targets, MEIOC-YTHDC2-RBM46 represses transcriptional repressors E2F6 and MGA, thereby allowing spermatogenic cells to activate *Meiosin* expression in response to retinoic acid. In turn, this activates the key meiotic transcriptional regulator STRA8-MEIOSIN, which amplifies expression of meiosis- and cell cycle-associated genes and drives the meiotic G1/S transition. In total, the post-transcriptional activity of MEIOC-YTHDC2-RBM46 enhances the activity of the meiotic transcriptional regulator. This regulatory pathway, acting in parallel with SCF complex-mediated degradation of DMRT1, enhances spermatogenic cells’ competence to initiate meiosis in response to retinoic acid.

## MATERIALS AND METHODS

### Animals

All experiments involving mice were performed in accordance with the guidelines of the Massachusetts Institute of Technology (MIT) Division of Comparative Medicine and Cincinnati Children’s Hospital Medical Center (CCHMC) Division of Veterinary Services, which are overseen by their respective Institutional Animal Care and Use Committees (IACUC). The animal care programs at MIT/Whitehead Institute and CCHMC are accredited by the Association for Assessment and Accreditation of Laboratory Animal Care, International (AAALAC) and meet or exceed the standards of AAALAC as detailed in the Guide for the Care and Use of Laboratory Animals. This research was approved by the MIT IACUC (no. 0617-059-20) and CCHMC IACUC (no. 2022-0061).

Mice carrying the *Meioc*-null allele *Meioc^tm1.1Dpc^*(Soh et al., 2017) were backcrossed to C57BL/6N (B6N) from Taconic Biosciences for at least 10 generations. Mice used for scRNA-seq experiments were also heterozygous for *Hspa2^tm1Dix^* (RRID: IMSR_JAX:013584;(Dix et al., 1996)) and homozygous for *Gt(ROSA)26Sor^tm9(CAG-tdTomato)Hze^*(*ROSA26^tdTomato^*; RRID:IMSR_JAX:007909; (Madisen et al., 2010)) with the floxed stop codon intact; both of these genotypes exhibit normal spermatogenesis.

### 10x Genomics single-cell RNA-seq

Single-cell sequencing libraries were prepared and sequenced in two batches, with each batch containing one wild-type and one *Meioc*-null pup at P15. One P15 testis per pup was enzymatically dissociated into single cells (see Supplementary Material and Methods) and resuspended in 0.05% bovine serum albumin (BSA) in phosphate buffered saline (PBS) for a target concentration of 1000 cells per microliter. Cell suspensions were loaded onto the Chromium Controller, aiming for recovery of 10,000 cells per sample. Libraries were generated using the Chromium Next GEM Single Cell 3ʹ v3.1 (10x Genomics), according to manufacturer’s instructions, and sequenced as 150-bp paired-end reads on an Illumina NovaSeq 6000 system with an S4 flow cell.

### Analysis of 10x Genomics scRNA-seq data

Alignment, filtering, barcode counting, and UMI counting were done via *count* function in Cell Ranger v.4.0.0 with default settings using Cell Ranger’s mm10-2020-A reference package (i.e., the GRCm38/mm10 mouse genome assembly with GENCODE vM23/Ensembl 98 annotation). Using Seurat v.3.2.3 (Stuart et al., 2019), cells were filtered for less than 10% mitochondrial reads, more than 1000 detected features, and a doublet score (generated by the *bcds* in scds v.1.2.0) of less than 0.4. Using protein-coding genes, UMI counts from both wild-type and *Meioc*-null samples were integrated and clusters identified using the first 30 dimensions.

The wild-type samples were used to assign cell types to clusters. Five somatic cell types were identified based on cell type-enriched gene expression: fetal Leydig cells (*Dlk1*, *Cyp11a1*) (Kaftanovskaya et al., 2015; Ye et al., 2017), peritubular myoid cells (*Acta2*, *Myh11*) (Chen et al., 2016; Cool et al., 2008), vascular endothelium (*Tm4sf1*) (Shih et al., 2009), testicular macrophages (*Cd14*, *Adgre1*, *Itgam*) (Bhushan et al., 2020; Landmann et al., 2000), and Sertoli cells (*Sox9*, *Cldn11*) (Mazaud-Guittot et al., 2010; Sekido et al., 2004). Germ cells were identified via *Ddx4* and *Dazl*. Then, using additional cell type-enriched gene expression along with UMAP-based cluster relationships, subclusters of spermatogenic cells were assigned to the following cell types: spermatogonial stem cells (*Id4*, *Gfra1*, *Etv5*) (He et al., 2007; Helsel et al., 2017; Sun et al., 2015; Wu et al., 2011), undifferentiated spermatogonia (*Zbtb16*) (Hobbs et al., 2012), type A spermatogonia (*Kit*, *Stra8*, *Ccnd2*) (Beumer et al., 2000; Endo et al., 2015; Schrans-Stassen et al., 1999; Yoshinaga et al., 1991), intermediate/type B spermatogonia (*Kit*) (Schrans-Stassen et al., 1999; Yoshinaga et al., 1991), preleptotene spermatocytes (*Stra8*), leptotene and zygotene spermatocytes (*Meiob*) (Chen et al., 2018; Souquet et al., 2013), and pachytene spermatocytes (*Piwil1*) (Chen et al., 2018; Deng and Lin, 2002).

Spermatogenic clusters were further refined based on cell cycle phase using the *CellCycleScoring* function in Seurat to identify type B spermatogonia in G2/M phase, as well as preleptotene spermatocytes in G1, early S, and late S phases (Fig. S1C,D). Clusters were merged as needed. Cell cycle designations of the preleptotene clusters were confirmed via gene expression patterns in wild-type cells independent of the gene:cell cycle phase pairings used by Seurat’s *CellCycleScoring* function (Fig. S3). These final cell type designations were then applied to the *Meioc*-null samples. The Mutant-only (Mut) cluster was primarily composed of *Meioc*-null spermatocyte cells, and only four wild-type germ cells were assigned to this cluster. The assignment of the 4 wild-type cells to the Mut cluster was considered to be a scRNA-seq artifact and these cells were excluded from subsequent analysis.

In addition to the somatic and germ cell clusters shown in Fig. S1A,B, one additional somatic cell cluster and one additional germ cell cluster were identified but could not be assigned a cell type using marker expression. This unassigned germ cell cluster is likely a technical artifact as it exhibits a median number of features lower than that of all other assigned germ cell subpopulations (Fig. S2A). The unassigned somatic and germ cell clusters, along with all other somatic clusters, were excluded from subsequent analyses. See Supplementary Materials and Methods for additional details on cluster-specific enrichment or depletion in wild-type cells and Gene Set Enrichment Analysis (GSEA).

Pseudotime trajectories were built in Monocle 3 v1.0.0 (Qiu et al., 2017a; Qiu et al., 2017b; Trapnell et al., 2014) using the following procedure: the Seurat object containing germ cell clusters only was imported as a Monocle object; data were normalized and pre-processed (function *preprocess_cds()*; options: num_dim = 100); batch effects were removed (function *align_cds()*; options: alignment_group = "batch"); dimensionality reduction was done via UMAP (function *reduce_dimension()*); cells were clustered (function *cluster_cells()*); a principal graph was learned from the reduced dimension space (function *learn_graph()*); and cells were ordered by selecting the Undiff cluster as the starting branch of the trajectory (function *order_cells()*). Cluster cell-type assignments made in Seurat were maintained in the pseudotime trajectories.

For differential expression analysis between wild-type and *Meioc*-null germ cells, log_2_ fold change was defined as wild-type over *Meioc*-null germ cells, such that the value reflects MEIOC’s activity in the unperturbed wild-type state. Differential expression analysis of scRNA-seq data was done on each germ cell cluster on genes with a minimum absolute log_2_ fold change of 0.1 and that were detected in at least 25% of either population using Seurat’s *FindMarkers* function (options: logfc.threshold = 0.1, min.pct = 0.25); *P* values were adjusted for multiple hypothesis testing of tested genes across all nine germ cell clusters via the Bonferroni correction using the *p.adjust* function in R. Additional expressed genes (i.e., genes detected in at least 25% of cells in wild-type or *Meioc*-null cells per cluster) that failed to meet the log_2_ fold change threshold for differential expression testing were identified via the *FindMarkers* function (options: logfc.threshold=0, min.pct = 0.25), and their *P* values were marked as “nd” for differential expression testing “not done.” See Supplementary Material and Methods for additional details on Gene Ontology analysis, cell cycle analysis of differentially expressed genes, and de novo motif analysis.

Dot plots were generated using Seurat’s *DotPlot* function with parameter scale=FALSE to maintain average expression from wild-type and *Meioc*-null samples on the same scale.

### Synchronization of spermatogenesis

Spermatogenesis was synchronized using a protocol originally developed by Hogarth et al. (Hogarth et al., 2013) and modified by Romer et al. (Romer et al., 2018). See Supplementary Material and Methods for details.

### Bulk RNA-seq analysis of preleptotene-enriched testes

Total RNA was extracted from four wild-type samples and three *Meioc*-null samples, using 1.5 synchronized testes from a single pup per sample. TRIzol Reagent (Thermo Fisher Scientific) was added to freshly thawed whole testes, and ERCC RNA ExFold RNA Spike-In Mix 1 and 2 (Thermo Fisher Scientific) were added to wild-type and *Meioc*-null samples, respectively, at a concentration of 1 μl of a 1:100 dilution of spike-in mix per 1 mg of testis tissue. (Spike-in mixes were ultimately not used for data analysis.) Total RNA was then isolated with chloroform following the manufacturer’s protocol, precipitated via isopropanol, and resuspended in RNase-free water. RNA-seq libraries were prepared with the TruSeq Stranded Total RNA kit with the Ribo-Zero Gold rRNA Removal kit. The barcoded libraries were pooled and sequenced with 50-bp single-end reads on an Illumina HiSeq 2500 machine.

Reads were quality trimmed using cutadapt v1.8 (options: -q 30 --minimum-length 20 -b AGATCGGAAGAGC). Expression levels of all transcripts in the mouse Gencode Basic vM15 gene annotation were estimated using kallisto v0.44.0 (Bray et al., 2016) with sequence-bias correction (--bias) and strand-specific pseudoalignment (--rf-stranded). Quantified transcripts were filtered for protein-coding genes, transcript-level estimated counts and transcripts per million (TPM) values were summed to the gene level, and TPMs were renormalized to transcript-per-million units.

To identify the MEIOC-dependent differential expression program, read counts from kallisto were rounded to the nearest integer and then supplied to DESeq2 v1.26.0 (Love et al., 2014). Genes were filtered for a minimum TPM of 1 in at least three of seven 2S preleptotene samples. Differential expression was defined using a cutoff of adjusted *P* value < 0.05. See Supplementary Material and Methods for additional details on cell cycle analysis, Gene Ontology analysis, and de novo motif analysis.

For analysis of transcript stability and transcriptional rates, reads were mapped to the mouse genome (mm10) with the GENCODE Basic vM15 gene annotation via STAR v2.7.1a (Dobin et al., 2013) (options: --outFilterMultimapNmax 1 --alignEndsType Extend5pOfRead1 -- outFilterMismatchNmax 2 --outSAMattributes None). All other parameters were set to default. Counts were quantified by htseq v0.11.0 (options: -m union --stranded=reverse) at the gene level (-type gene) and exon level (-type exon), and intron levels were calculated as gene-level counts minus exon-level counts. Changes in transcript stability, transcriptional rate, and abundance were calculated for each gene for each sample using REMBRANDTS (Alkallas et al., 2017) with a stringency of 0.80 and linear bias mode.

The cumulative distribution of the log_2_ fold change (WT/*Meioc* KO) in transcript stability or transcriptional rate in MEIOC-upregulated or -downregulated transcripts was compared to expressed but unchanged transcripts via a two-tailed Wilcoxon rank sum test with Bonferroni correction for multiple hypothesis testing using the *wilcox.test* and *p.adjust* functions in R. MEIOC-upregulated genes were defined as log_2_ fold change WT/*Meioc* KO > 0 and *P* < 0.05 and MEIOC-downregulated genes were defined as log_2_ fold change WT/*Meioc* KO < 0 and *P* < 0.05 based on the DESeq2 analysis of the bulk RNA-seq data.

### Re-analysis of STRA8 RNA-seq dataset

RNA-seq data from *Stra8*-null and *Stra8*-heterozygote (i.e., phenotypically wild-type) preleptotene spermatocytes (NCBI GEO GSE115928; Kojima et al., 2019) were reanalyzed. Genes with STRA8-bound promoters identified by ChIP-seq in testes enriched for preleptotene spermatocytes were extracted from Supplementary File 2 (Kojima et al. 2019) (https://doi.org/10.7554/eLife.43738.028). See Supplementary Material and Methods for details.

### Re-analysis of MEIOC RIP-seq dataset and comparison to YTHDC2 and RBM46 CLIP datasets

MEIOC RIP-seq from P15 testes (NCBI GEO GSE96920; Soh et al., 2017) were reanalyzed, with additional details in Supplementary Material and Methods.

YTHDC2-bound mRNAs identified via CLIP-seq in P8 and P10 testes were extracted from Tables S1 and S2 (Saito et al., 2022) (http://genesdev.cshlp.org/content/suppl/2022/01/19/gad.349190.121.DC1/Supplemental_Tables.xlsx).

RBM46 eCLIP data from P12-P14 testes (NCBI GEO GSE197282) (Qian et al., 2022) were reanalyzed, with additional details in Supplementary Material and Methods.

### Identification of transcriptional repressors that are repressed by MEIOC-YTHDC2-RBM46

To identify transcriptional repressors that are repressed by MEIOC-YTHDC2-RBM46, we first identified transcripts that were (i) downregulated by MEIOC in the In/B, B G2M, and/or pL G1 clusters and (ii) targeted by MEIOC-YTHDC2-RBM46. We then manually examined the “Function” description of the UniProtKB database (release 2020_01; www.uniprot.org) for these mouse proteins. Any proteins that were annotated as repressing transcription or gene expression were considered transcriptional repressors.

### Re-analysis of E2F6 and MGA ChIP-seq datasets and RNA-seq datasets from ESCs

The following datasets were reanalyzed here: E2F6 ChIP-seq and input data from wild-type and *E2f6*-knockout mouse ESCs (NCBI GEO GSE149025) (Dahlet et al., 2021); MGA and IgG ChIP-seq from wild-type mouse ESCs (ArrayExpress E-MTAB-6007) (Stielow et al., 2018); mouse ESC RNA-seq data from wild-type and *E2f6*-knockout samples (NCBI GEO GSE149025) (Dahlet et al., 2021); and wild-type and *Mga*-knockout samples (NCBI GEO GSE144141) (Qin et al., 2021). See supplementary Material and Methods for details.

### Immunostaining

Immunostaining was carried out in tissue sections using the following primary antibodies: anti-DMRT1 (Santa Cruz Biotechnology sc-377167, 1:200 dilution for fluorescent staining); anti-MEIOSIN (guinea pig polyclonal from Ishiguro et al., 2020, 1:100 dilution for fluorescent staining); anti-STRA8 (Abcam ab49405, 1:500 dilution for chromogenic staining or 1:200 for fluorescent staining); and anti-STRA8 (Abcam ab49602, 1:200 dilution for fluorescent staining). See supplementary Material and Methods for details.

## Supporting information

Supplemental methods, references, and figures

## Acknowledgements

We thank Kei-Ichiro Ishiguro for the MEIOSIN antibody; Mary Goodheart, Mina Kojima, Holly Christensen, and Peter Nicholls for technical help; and Abigail Groff and Linyong Mao for advice on bioinformatics analysis. We are grateful to the members of the Page Lab for valuable feedback, and to Winston Bellott, Jordana Bloom, ZY Chen, Jennifer Hughes, Mina Kojima, and Adrianna San Roman for comments on the manuscript.

We thank the Whitehead Genome Technology Core for library preparation and Illumina sequencing, and the Whitehead Institute FACS Facility for cell sorting. Fluorescent microscopy was done at the Whitehead Institute’s WM Keck Foundation Biological Imaging Facility and Cincinnati Children’s Bio-Imaging and Analysis Facility (RRID:SCR_022628).

## Author Contributions

M.M.M and B.L. prepared samples for 10x Genomic analysis. D.deR. assessed germ cell stage in synchronized testis samples. M.M.M. and N.G.P. carried out immunostaining experiments.

M.M.M. performed all other experiments and data analysis. M.M.M. and D.C.P. designed the study and wrote the manuscript with contributions from all authors.

## Data availability

Sequencing data have been deposited to the Gene Expression Omnibus under accession number GSE230747.

