## Supplemental methods, references, and figures for "Destabilization of mRNAs enhances competence to initiate meiosis in mouse spermatogenic cells"

#### SUPPLEMENTARY MATERIALS AND METHODS

*Enzymatic digestion of postnatal testis in preparation for 10x Genomics single-cell RNA-seq*

One P15 testes per sample was enzymatically dissociated by shaking vigorously for 30 sec and incubating at 35˚C for 7 min in 900 ul of 0.1% collagenase type I (Worthington LS004196) in Hank's Balanced Salt Solution (HBSS), supplemented with TURBO DNase (Thermo Fisher Scientific AM2238) for a final concentration of 2U $mL^{-1}$. After allowing the sample to sit for 1 min such that the seminiferous tubules settled to the bottom of the tube, the interstitium was removed by discarding the resulting supernatant, and the remaining sample was incubated at 35˚C for 25 min (with gently pipetting every 5 minutes) in 800 ul of 0.05% trypsin and 0.1% collagenase type I in HBSS, supplemented with TURBO DNase to a final concentration of 2U $mL^{-1}$. The cell suspension was then passed through a 40 um nylon strainer, and cell suspension was allowed to digest at 35˚C for an additional 20 min. The digestion was quenched by adding 200ul fetal bovine serum (FBS) and gently mixing. The cell suspension was again passed through a 40 um nylon strainer. The concentration of the resulting cell suspension was assessed using trypan blue on the Countess Cell Counter. Cells were pelleted at 500g for 4 min at room temperature, resuspended in 0.05% bovine serum albumin (BSA) in phosphate buffered saline (PBS) for a target concentration of 1000 cells per microliter, and placed on ice. Viability and concentration were re-assessed ~20 min prior to loading on the Chromium Controller using trypan blue on a Countess Cell Counter, at which point all samples had a minimum of 90% viability.

*Identification of cell type-specific markers and Gene Set Enrichment Analysis in scRNA-seq data*

Using wild-type samples only, markers that were depleted or enriched within a cell type were identified via Seurat's *FindAllMarkers* function without a fold change threshold (options: only.pos = FALSE, logfc.threshold = 0, min.pct = 0.25). Features were scaled and centered via Seurat's *ScaleData* function, using percent mitochondrial reads and batch as variables that were regressed out, before generating a heatmap via the *DoHeatmap* function. Gene Set Enrichment Analysis (GSEA) was done using all genes that were statistically tested within each cluster, with $P$ values of 0 replaced with $10^{-300}$ (i.e., smaller than the smallest non-zero $P$ value) and ranks based on the signed $-\log_{10} P$ value. GSEA was done on Gene Ontology (GO) terms that contained between 100 and 500 genes via clusterProfiler v3.0.4's *gseGO* function (options: OrgDb = "org.Mm.eg.db", ont = "ALL", minGSSize = 100, maxGSSize = 500, pvalueCutoff = 0.05), and the three GO terms exhibiting the greatest enrichment based on Normalized Enrichment Score (NES) were graphed via enrichplot v1.16.2's *gseaplot2* function in R (Yu et al., 2012).

*Synchronization of spermatogenesis*

Spermatogenesis was synchronized using a protocol originally developed by Hogarth et al. (Hogarth et al., 2013) and modified by Romer et al. (Romer et al., 2018). Briefly, male mice were injected daily subcutaneously from postnatal day (P) 2 to P8 with WIN 18,446 (Santa Cruz Biotechnology) at 0.1mg $g^{-1}$ body weight and on P9 with retinoic acid (RA; MilliporeSigma) at 0.0125 mg $g^{-1}$ body weight. To obtain testes enriched for preleptotene cells, mice were euthanized and testes were collected 6.75 d after the RA injection. For each pup, a small testis biopsy was collected for histology to confirm proper enrichment of the desired cell type, and the rest of the testes were flash frozen as whole testes or cell sorted. For histological verification of staging, the testis biopsy was fixed in Bouin's solution and stained for STRA8, as described below. For each pair of testes, at least 40 tubule cross-sections were staged based on the most advanced spermatogenic cell stage present. All samples analyzed via bulk RNA-seq contained at least 87.5% of tubule cross-sections in the preleptotene stage.

*Gene Ontology analysis of scRNA-seq and bulk RNA-seq data*

Within each scRNA-seq cluster, MEIOC-upregulated ($\log_2$ fold change > 0.1 and adjusted *P* value < 0.05) and MEIOC-downregulated genes ($\log_2$ fold change < -0.1 and adjusted *P* value < 0.05) were separately tested for enrichment of Gene Ontology Biological Processes gene lists relative to a background of all expressed genes (i.e., genes expressed in at least 25% of wild-type or *Meioc*-null cells within that cluster). For bulk RNA-seq data, MEIOC-upregulated ($\log_2$ fold change > 0 and adjusted *P* value < 0.05) and MEIOC-downregulated genes ($\log_2$ fold change < -0.1 and adjusted *P* value < 0.05) were similarly tested against a background of all expressed genes (i.e., a minimum TPM of 1 in at least three of seven 2S preleptotene samples). These analyses were done via R package clusterProfiler v3.0.4 using the *enrichGO* function (options: OrgDb = "org.Mm.eg.db", ont = "BP", pvalueCutoff = 0.05, readable = T, pAdjustMethod =

"BH"). For the scRNA-seq data, the average $log_2$ fold change for select upregulated and downregulated genes that fell under GO terms “meiotic cell cycle” (GO:0051321) and “mitotic cell cycle” (GO:0000278), respectively, were graphed via heatmap to show gene expression changes between wild-type and *Meioc*-null cells across scRNA-seq germ cell clusters.

*Cell cycle analysis of bulk RNA-seq and differentially expressed genes from scRNA-seq’s late S phase preleptotene (pL lS), leptotene (L), and zygotene (Z) clusters*

A curated list of human cell cycle genes whose expression is associated with specific cell cycle phases was obtained from Supplementary File S1 of Hsiao et al., 2020. This gene set originated from Macosko et al., 2015 and represents a subset of the genes annotated in Whitfield et al., 2002. (Note that this Macosko et al., 2015 gene list was also used for the cell cycle analysis implemented for scRNA-seq analysis by Seurat.) These human genes were converted to their one-to-one mouse orthologs, resulting in a list of 564 mouse genes.

For analysis of bulk RNA-seq data, percentile ranks were calculated for all expressed genes using the $log_2$ fold changes (WT/*Meioc* KO) from the differential expression results with 0 representing the most downregulated genes and 1 representing the most upregulated genes. Genes’ percentile ranks for each cell cycle phase was compared to the mean percentile rank for all expressed genes (0.5) via a two-sided Wilcoxon signed rank test with Bonferroni correction for multiple hypothesis testing, as implemented by the *wilcox.test* and *p.adjust* functions in R.

To assess what cell cycle phase was represented by the enrichment for the “mitotic cell cycle” GO term (GO:0000278) in MEIOC-downregulated genes among the pL, L, and Z clusters, the downregulated genes associated with this GO term were extracted and linked to their associated cell cycle phase (if any), as annotated by Hsiao et al., 2020. The percentage of these genes associated with G1/S, S, G2, M, and M/G1 were graphed. Note that some genes are associated with more than one cell cycle phase and are therefore represented more than once in the graph. Genes whose expression was not associated with a specific cell cycle phase were not graphed.

*Re-analysis of MEIOC RIP-seq dataset and comparison to YTHDC2 and RBM46 CLIP datasets*

MEIOC RIP-seq from P15 testes (NCBI GEO GSE96920; Soh et al., 2017) was reanalyzed. Reads were quality trimmed using the methods described above. For the MEIOC RIP-seq analysis, the additional cutadapt option --cut 3 was used to remove each sequencing read's first three bases that were added during SMARTer Stranded library preparation. Trimmed reads were pseudoaligned using the methods described above, with the option --fr-stranded for strand-specificity. Quantified transcripts were filtered for protein-coding genes, transcript-level estimated counts and transcripts per million (TPM) values were summed to the gene level, and TPMs were renormalized to transcript-per-million units. Read counts from kallisto were rounded to the nearest integer and then supplied to DESeq2 v1.26.0.Targets for each RIP-seq dataset were defined using a DESeq model as previously described (Soh et al., 2017), using cutoffs of fold change > 1.5 and adjusted *P* value < 0.05. (This fold change cut off was less stringent than our prior published analysis, which used fold change >3 (Soh et al., 2017).) MEIOC targets and nontargets were limited to those genes with a minimum TPM of 1 in at least 2 of 4 wild-type input RNA-seq samples.

RBM46 eCLIP data from P12-P14 testes (NCBI GEO GSE197282) (Qian et al., 2022) was reanalyzed using Read 1 fastq files only. CLIP data was analyzed as previously described (Busch et al., 2020): sequencing reads were mapped to the mouse genome (mm10) with the GENCODE Basic vM15 gene annotation via STAR v2.7.1a (options: --outFilterMismatchNoverReadLmax 0.04 --outFilterMultimapNmax 1 --alignEndsType Extend5pOfRead1 --outFilterMismatchNmax 2 --outSJfilterReads Unique), and peaks were called via PureCLIP v1.3.1 (Krakau et al., 2017) (options: -nt 8 -ld -iv 'chr1;chr2;chr3;') using two iCLIP biological replicates and one input. Peaks were filtered for those that were supported by both eCLIP biological replicates, annotated using the GENCODE Basic vM15 gene annotation, and assigned to a transcript position based on the following hierarchy: 3' UTR exon > 5' UTR exon> coding exon > intron. In total, we identified 19,667 peaks corresponding to 4,912 protein-coding genes, which is similar to the previously reported 24,010 peaks and 4,413 protein-coding genes (Qian et al., 2022).

The overlaps between MEIOC targets and YTHDC2 targets; MEIOC targets and RBM46 targets; as well as MEIOC-YTHDC2 targets and RBM46 targets were tested for enrichment via one-tailed hypergeometric tests with Bonferroni correction for multiple hypothesis testing using the *phyper* (options: lower.tail = *F* ) and *p.adjust* functions in R.

For Gene Ontology analysis of MEIOC-YTHDC2-RBM46 targets and MEIOC-RBM46 targets, each gene list was tested for enrichment of Gene Ontology Biological Processes gene lists relative to a background of all expressed genes in the MEIOC RIP-seq analysis (minimum TPM of 1 in at least 2 of 4 wild-type input RNA-seq samples) via R package clusterProfiler v3.0.4 using the *enrichGO* function (options: OrgDb = "org.Mm.eg.db", ont = "BP", pvalueCutoff = 0.05, readable = T, pAdjustMethod = "BH").

*Comparison of MEIOC, YTHDC2, and RBM46 targets to MEIOC scRNA-seq and bulk RNA-seq datasets*

The overlaps between bound mRNAs and MEIOC-upregulated or -downregulated genes were tested for enrichment via one-tailed hypergeometric tests with Bonferroni correction for multiple hypothesis testing using the *phyper* (options: lower.tail = *F* ) and *p.adjust* functions in R. For scRNA-seq, MEIOC-upregulated genes were defined as $\log_2$ fold change WT/*Meioc* KO > 0.1 and $P < 0.05$; MEIOC-downregulated genes were defined as $\log_2$ fold change WT/*Meioc* KO < -0.1 and $P < 0.05$. For bulk RNA-seq, MEIOC-upregulated genes were defined as $\log_2$ fold change WT/*Meioc* KO > 0 and $P < 0.05$; MEIOC-downregulated genes were defined as $\log_2$ fold change WT/*Meioc* KO < 0 and $P < 0.05$.

The cumulative distribution of the $\log_2$ fold change (WT/*Meioc* KO) per cluster-specific scRNA-seq dataset for all MEIOC targets was compared to nontargets via a two-tailed Wilcoxon rank sum test with Bonferroni correction for multiple hypothesis testing using the *wilcox.test* and *p.adjust* functions in R. A similar approach was used to compare the cumulative distribution of MEIOC-YTHDC2-RBM46 targets, MEIOC-RBM46-only targets, and MEIOC-YTHDC2-only targets to MEIOC-only targets as well as MEIOC-only to nontargets.

The overlaps between MEIOC-YTHDC2-RBM46 targets and GO term “mitotic cell cycle genes” downregulated by MEIOC as well as MEIOC-YTHDC2-RBM46 targets and GO term “meiotic cell cycle genes” upregulated by MEIOC were tested for enrichment via one-tailed hypergeometric tests with Bonferroni correction for multiple hypothesis testing using the *phyper* (options: lower.tail = *F* ) and *p.adjust* functions in R. For scRNA-seq, MEIOC-upregulated genes were defined as $\log_2$ fold change WT/*Meioc* KO > 0.1 and $P < 0.05$ for any cluster; MEIOC-downregulated genes were defined as $\log_2$ fold change WT/*Meioc* KO < -0.1 and $P < 0.05$ for any cluster. For bulk RNA-seq, MEIOC-upregulated genes were defined as $\log_2$ fold change WT/*Meioc* KO > 0 and $P < 0.05$; MEIOC-downregulated genes were defined as $\log_2$ fold change WT/*Meioc* KO < 0 and $P < 0.05$.

The cumulative distribution of the $\log_2$ fold change (WT/*Meioc* KO) in transcript stability (defined by REMBRANDTS analysis of bulk RNA-seq data) for MEIOC targets was compared to that for nontargets via a two-tailed Wilcoxon rank sum test using the *wilcox.test* function in R. A similar approach was used to compare the cumulative distribution of MEIOC-YTHDC2-RBM46 targets, MEIOC-RBM46-only targets, and MEIOC-YTHDC2-only targets to MEIOC-only targets as well as MEIOC-only targets to nontargets, with the addition of Bonferroni correction for multiple hypothesis testing using the *p.adjust* function in R.

*Re-analysis of E2F6 and MGA ChIP-seq datasets from ESCs*

E2F6 ChIP-seq and input data from wild-type and *E2f6*-knockout mouse ESCs (NCBI GEO GSE149025) (Dahlet et al., 2021) as well as MGA and IgG ChIP-seq from wild-type mouse ESCs (ArrayExpress E-MTAB-6007) (Stielow et al., 2018) were analyzed. Reads were quality trimmed as described above and mapped to the mouse genome (GRCm38/mm10 assembly) via bowtie2 v.2.3.4.1 (Langmead and Salzberg, 2012). ChIP-seq peaks were called via mac2 v.2.2.7.1 (Zhang et al., 2008) with a default $P$ value cutoff of $1\times10^{-5}$. E2F6 ChIP-seq peaks were called using the corresponding input sample, while MGA ChIP-seq peaks were called using the IgG ChIP-seq sample as the control. Peaks were annotated using annotatePeaks.pl from the HOMER motif discovery software v.4.11.1 (Heinz et al., 2010) with the mm10 genome v.6.3, and those overlapping gene promoters (defined as transcriptional start site ±1000bp) were identified. Peaks identified in *E2f6*-knockout cells were considered background signal from the antibody, and peaks associated with a gene promoter in both the wild-type and *E2f6*-knockout cells were excluded from subsequent analysis, regardless of the extent of peak overlap between the two samples.

Input-subtracted normalized ChIP-seq signal was visualized by generating a bigWig file with the input chromatin reads subtracted from the ChIP sample reads using the bamCompare from deepTools v.3.5.0 (Ramírez et al., 2016) (options: --operation subtract --scaleFactorsMethod None --normalizeUsing RPKM --ignoreDuplicates --smoothLength 100 --binSize 10). For E2F6 ChIP-seq, input samples were used as input; for MGA ChIP-seq, the IgG ChIP-seq sample was used as input. ChIP-seq signal was visualized using the GRCm38/mm10 genome assembly on the UCSC Genome Browser (http://genome.ucsc.edu/).

Input-subtracted ChIP-seq signal at gene promoters was quantified by first removing potential PCR duplicates using rmdup from samtools v.1.11 (Li et al., 2009). The number of reads at the promoter (transcriptional start site ±1000bp) of each protein-coding transcript from the GENCODE Basic vM15 annotation was determined and summarized on a per-gene basis using htseq-count (options: -m union -t transcript -i gene_name) (Anders et al., 2015). For each gene, counts were normalized to reads per million (rpm) by scaling to the number of unduplicated reads, and the normalized input read count was subtracted from the normalized ChIP-seq-read count.

*Re-analysis of RNA-seq datasets from wild-type,* E2f6*-knockout, and* Mga*-knockout ESCs*

Mouse ESC RNA-seq data from wild-type and *E2f6*-knockout samples (NCBI GEO GSE149025) (Dahlet et al., 2021) as well as from wild-type and *Mga*-knockout samples (NCBI GEO GSE144141) (Qin et al., 2021) were analyzed. Reads were quality trimmed and pseudoaligned as described above, except without the strand-specific pseudoalignment option. Quantified transcripts were filtered for protein-coding genes, transcript-level estimated counts and transcripts per million (TPM) values were summed to the gene level, and TPMs were renormalized to transcript-per-million units. Read counts from kallisto were rounded to the nearest integer and then supplied to DESeq2 v1.26.0. Within each dataset, differential expression was determined using a DESeq model that included genotype. Genes were filtered for a minimum TPM of 1 in at least two of six samples. For differential expression analysis, $\log_2$ fold change was defined as wild-type over *E2f6* or *Mga*-null germ cells, such that the values reflects E2F6 or MGA's activity in the unperturbed wild-type state.

E2F6-activated and -repressed genes were defined as (i) E2F6-upregulated ($\log_2$ fold change WT/*E2f6* KO > 0 and $P < 0.05$) and -downregulated ($\log_2$ fold change WT/*E2f6* KO < 0 and $P < 0.05$) genes, respectively, and (ii) E2F6-bound promoters from E2F6 ChIP-seq reanalysis. MGA-activated and -repressed genes were similarly defined. The overlaps between E2F6-activated and MGA-activated genes; E2F6-activated and MGA-repressed genes; E2F6-repressed and MGA-activated genes; and E2F6-repressed and MGA-repressed genes were tested for enrichment via one-tailed hypergeometric tests with Bonferroni correction for multiple hypothesis testing using the *phyper* (options: lower.tail = *F* ) and *p.adjust* functions in R.

*Comparison of genes regulated by E2F6 and MGA to MEIOC scRNA-seq and bulk RNA-seq datasets*

The overlaps between E2F6- and/or MGA-regulated genes and MEIOC-upregulated or -downregulated genes were tested for enrichment via one-tailed hypergeometric tests with Bonferroni correction for multiple hypothesis testing using the *phyper* (options: lower.tail = *F* ) and *p.adjust* functions in R. For scRNA-seq, MEIOC-upregulated genes were defined as $\log_2$ fold change WT/*Meioc* KO > 0.1 and $P < 0.05$; MEIOC-downregulated genes were defined as $\log_2$ fold change WT/*Meioc* KO < -0.1 and $P < 0.05$. For bulk RNA-seq, MEIOC-upregulated genes were defined as $\log_2$ fold change WT/*Meioc* KO > 0 and $P < 0.05$; MEIOC-downregulated genes were defined as $\log_2$ fold change WT/*Meioc* KO < 0 and $P < 0.05$.

The overlap between E2F6 and/or MGA-repressed genes and GO term "meiotic cell cycle" genes upregulated by MEIOC ($\log_2$ fold change WT/*Meioc* KO > 0.1 and $P < 0.05$) in the In/B, B G2/M, or pL G1 clusters was tested for enrichment via a one-tailed hypergeometric test using the *phyper* (options: lower.tail = *F* ) in R. A similar analysis was also done for genes upregulated by MEIOC as defined in the bulk RNA-seq analysis ($\log_2$ fold change WT/*Meioc* KO > 1 and $P < 0.05$).

*Comparison to RNA-seq dataset from* Stra8*-null preleptotene spermatocytes*

RNA-seq data from *Stra8*-null preleptotene spermatocytes (N=3) and *Stra8*-heterozygote preleptotene spermatocytes expressing high levels of *Stra8* (i.e., phenotypically wild-type controls; N=2), obtained via synchronization and sorting (NCBI GEO GSE115928; Kojima et al., 2019), was quality filtered, pseudoaligned, and analyzed for differential expression as described above. To determine whether genes upregulated by MEIOC were enriched for genes also upregulated by STRA8; whether genes downregulated by MEIOC were enriched for genes downregulated by STRA8; and whether genes upregulated by MEIOC were enriched for genes directly activated by STRA8, these gene lists were statistically compared using one-tailed hypergeometric tests with Bonferroni correction for multiple hypothesis testing via the *phyper* (options: lower.tail = *F*) and *p.adjust* functions in R.

For each scRNA-seq cluster, Spearman rank correlation rho ($\rho$) between the MEIOC scRNA-seq and STRA8 bulk RNA-seq datasets' $\log_2$ fold change for STRA8-dependent genes only (padj<0.05) was calculated using the *cor.test* function in R (options: method='spearman').

In the graphs displaying the MEIOC scRNA-seq and STRA8 bulk RNA-seq datasets' $\log_2$ fold change, the small number of extreme data points that fell outside of graph's axes are displayed as 0.1 outside of the axis limit. The same approach was taken to compare the MEIOC bulk RNA-seq dataset and the STRA8 bulk RNA-seq dataset.

For each scRNA-seq cluster, the bootstrapping analysis of each comparison of the MEIOC scRNA-seq and STRA8 bulk RNA-seq datasets' $\log_2$ fold change was done in R. Spearman rank correlation rhos were calculated for 100,000 gene sets randomly generated from all expressed genes in the two datasets. The number of sampled genes was equal to the number of STRA8-dependent genes in the two datasets. The Spearman rhos from random sampling were used to calculate a Z-score for the Spearman rho of the STRA8-dependent genes. *P*-value was calculated from the Z-score using the *pnorm* function in R (options: lower.tail = FALSE, log.p = TRUE). The same approach was taken to calculate a *P*-value to the Spearman rho between the MEIOC bulk RNA-seq and STRA8 bulk RNA-seq datasets.

STRA8-activated genes were defined as STRA8-upregulated genes ($\log_2$ fold change WT/*Stra8* KO > 0 and $P < 0.05$) with STRA8-bound promoters (as identified in Kojima et al., 2019). The overlaps between STRA8-activated genes and MEIOC-upregulated genes were tested for enrichment via one-tailed hypergeometric tests with Bonferroni correction for multiple hypothesis testing using the *phyper* (options: lower.tail = *F* ) and *p.adjust* functions in R. This approach was also used to test for enrichment between STRA8-activated genes repressed by E2F6 or MGA and MEIOC-upregulated genes; STRA8-activated genes not repressed by E2F6 or MGA and MEIOC-upregulated genes; STRA8-activated genes and E2F6- and MGA-repressed genes; STRA8-activated genes and genes repressed by E2F6 only; and STRA8-activated genes and genes repressed by MGA only. MEIOC-upregulated genes were defined as $\log_2$ fold change WT/*Meioc* KO > 0.1 and $P < 0.05$ in the scRNA-seq analysis and as $\log_2$ fold change WT/*Meioc* KO > 0 and $P < 0.05$ in the bulk RNA-seq analysis.

*Motif analysis of MEIOC-dependent genes from scRNA-seq and bulk RNA-seq data*

De novo motif analysis of the promoters (transcriptional start site ±1000 bp) of MEIOC-upregulated genes with the promoters of all expressed genes as background was performed via HOMER motif discovery software v.4.11.1 (Heinz et al., 2010) (options: -len 8,10) with the mm10 genome v.6.3. For scRNA-seq, de novo motif analysis was carried out for pL eS, pL lS, and L clusters, but all enriched motifs identified were likely false positives ($P$ >= 1E-10), as designated by HOMER. For bulk RNA-seq analysis, the top two statistically enriched motifs are reported in Fig. S12D.

*Chromogenic and fluorescent immunostaining*

For chromogenic staining of synchronized testes, testis biopsies were fixed in Bouin's solution for 2 hours at room temperature. For fluorescent staining, dissected testes were fixed in 4% (w $v^{-1}$) paraformaldehyde at 4˚C overnight. Fixed tissues were embedded in paraffin, and 6 um sections were generated. Slides were deparaffinized and hydrated using xylene and ethanol solutions, respectively. For antigen retrieval, slides were boiled in citrate buffer (10 mM sodium citrate, 0.05% Tween 20, pH 6.0) for 10 min.

For chromogenic staining, tissue sections were stained with anti-STRA8 (Abcam ab49405, 1:500 dilution) using ImmPRESS HRP anti-Rabbit Detection Kit (Vector Laboratories MP-7401–50) and ImmPACT DAB Peroxidase Substrate (Vector Laboratories SK-4105). Sections were then washed in PBS, counterstained with hematoxylin, and coverslipped with Permount Mounting Medium.

For fluorescent staining, tissue sections were incubated in blocking solution (10% normal donkey serum in PBS) for 1 hour at room temperature and then with primary antibodies diluted in blocking solution overnight at 4˚C. Sections were washed in PBS before incubating with fluorophore-conjugated secondary antibodies in blocking solution for 1 hour at room temperature. Sections were washed in PBS and counterstained with DAPI.

For quantification of MEIOSIN signal intensity in STRA8-positive cells from wild-type (*Meioc* +/+) and *Meioc* KO testes, P10 testis tissue sections from three wild-type and three *Meioc*-null animals (representing 3 wild-type:*Meioc*-null littermate pairs) were immunostained for STRA8 (Abcam ab49405, 1:200 dilution) and MEIOSIN (guinea pig polyclonal from Ishiguro et al., 2020, 1:100 dilution). The following secondary antibodies were used: donkey anti-guinea pig DyLight 647 (Jackson ImmunoResearch Laboratories 706-605-148, 1:250 dilution) and donkey anti-rabbit AlexaFluor 488 (Jackson ImmunoResearch Laboratories 711-545-152, 1:250 dilution). After counterstaining with DAPI, slides were coverslipped with ProLong Gold Antifade reagent (Thermo Fisher Scientific), and imaged via Zeiss LSM 700 laser scanning confocal microscope with Zeiss Plan Apochromat 40x/1.3 oil objective. Confocal images were acquired at a resolution of 4.8177 pixels per um. CellProfiler v4.0.7 was used to quantify MEIOSIN signal intensity in STRA8-positive and STRA8-negative cells. CellProfiler output was imported into R, and STRA8-positive cells were filtered for a minimum area (AreaShape_Area≥500 pixels). STRA8-positive cells were also filtered for a circular shape (1.3≥AreaShape_FormFactor≥0.7) to exclude STRA8-positive spermatogonia and limit analysis to round STRA8-positive preleptotene spermatocytes. For each image, background MEIOSIN signal was defined as the median MEIOSIN intensity in all STRA8-negative cells; then the background-subtracted median MEIOSIN intensity was calculated for each of the filtered STRA8-positive cells in that image. The STRA8-positive cells were then randomly downsampled to 75 cells per animal to ensure that all biological replicates were equally represented (Table S13). The background-subtracted median MEIOSIN intensity in STRA8-positive cells from wild-type and *Meioc*-null animals was compared via a two-sided Wilcoxon signed rank test, as implemented by the *wilcox.test* function in R.

For DMRT1 immunostaining, P15 testis tissue sections from two phenotypically wild-type (*Meioc* +/+ or +/-) and three *Meioc* KO animals from two litters were examined. These animals were also heterozygous for *Hspa2*$^{tm1Dix}$, a genotype that exhibits normal spermatogenesis. For antigen retrieval, slides were boiled in an alternative buffer (10 mM Tris, 1mM EDTA, 0.05% Tween 20, pH 9.0) for 12 min. Sections were immunostained for STRA8 (Abcam ab49602, 1:200 dilution) and DMRT1 (Santa Cruz Biotechnology sc-377167, 1:200 dilution). The following secondary antibodies were used: donkey anti-rabbit Alexa Fluor 568 (Abcam ab175692, 1:250 dilution) and donkey anti-mouse Alexa Fluor 488 (Invitrogen A-21202, 1:250 dilution). After counterstaining with DAPI, slides were coverslipped with ProLong Gold Antifade reagent (Thermo Fisher Scientific) and imaged via Nikon 90i upright widefield microscope.

Figure S1



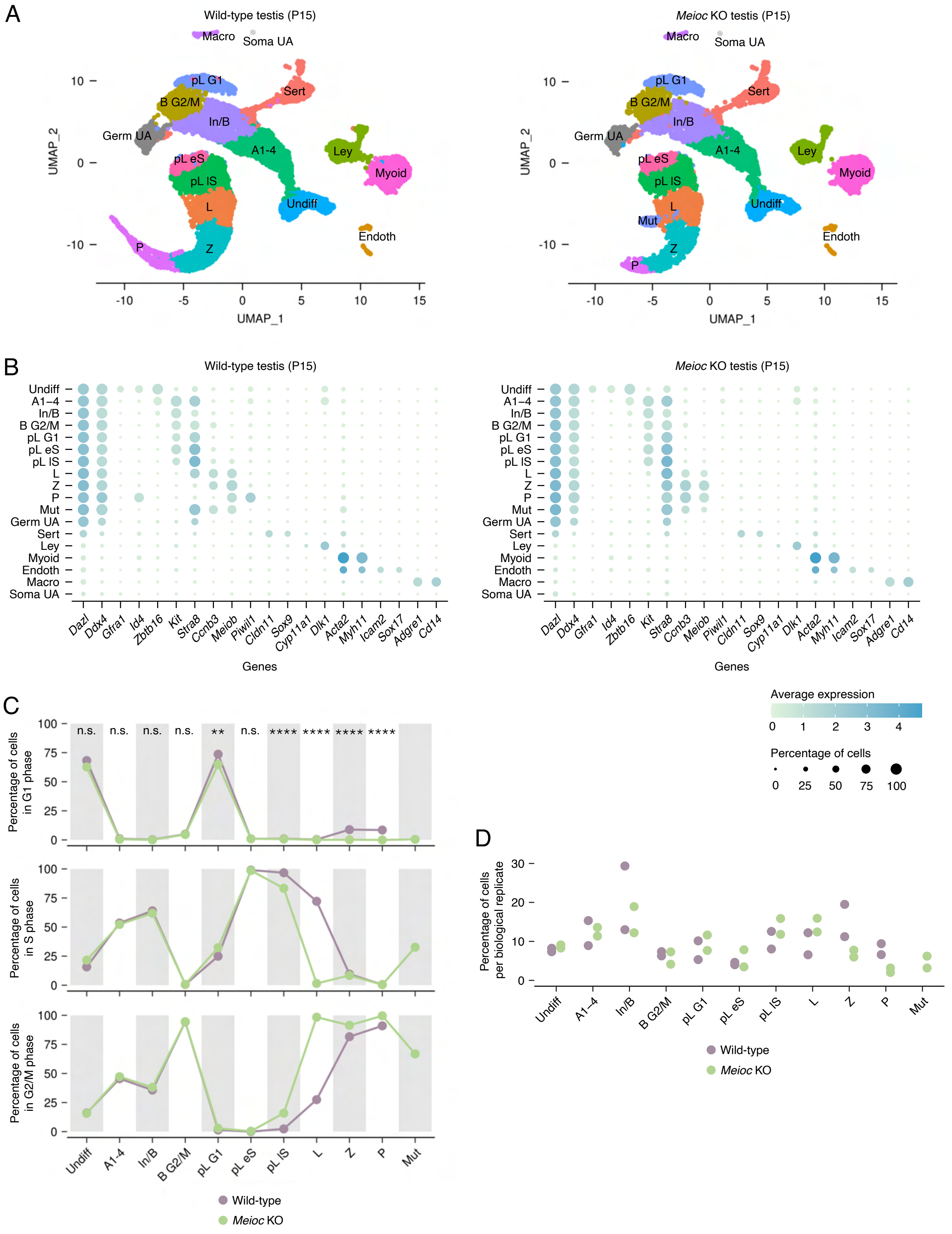
A
Wild-type testis (P15)
Meioc KO testis (P15)
UMAP_1
UMAP_2
Macro
Soma UA
pL G1
Sert
B G2/M
In/B
Germ UA
A1-4
Ley
pL eS
Myoid
pL IS
L
Undiff
Endoth
Z
P
Mut
B
Genes
Average expression
Percentage of cells
Dazl
Ddx4
Gfra1
Id4
Zbtb16
Kit
Stra8
Ccnb3
Meiob
Piwil1
Cldn11
Sox9
Cyp11a1
Dlk1
Acta2
Myh11
Icam2
Sox17
Adgre1
Cd14
C
Percentage of cells in G1 phase
Percentage of cells in S phase
Percentage of cells in G2/M phase
n.s.
**
****
Wild-type
Meioc KO
D
Percentage of cells per biological replicate

**Figure S1: Identification of germ cell subpopulations across the mitosis-to-meiosis transition via scRNA-seq.**

**A:** UMAP of all clusters identified via Seurat from wild-type and *Meioc* KO P15 testes (N=2 per genotype).
**B:** Dotplot of expression levels and percentage of cells for markers used to assign clusters.
**C:** Distribution of cell cycle phase per germ cell cluster from wild-type and *Meioc* KO samples.
**D:** Percentage of cells per cluster by biological replicate, relative to the total number of germ cells in these clusters.

*, adj. $P<0.05$; **, adj. $P<0.01$; ***, adj. $P<0.001$; adj. $P<0.0001$; n.s., not significant.

Figure S2



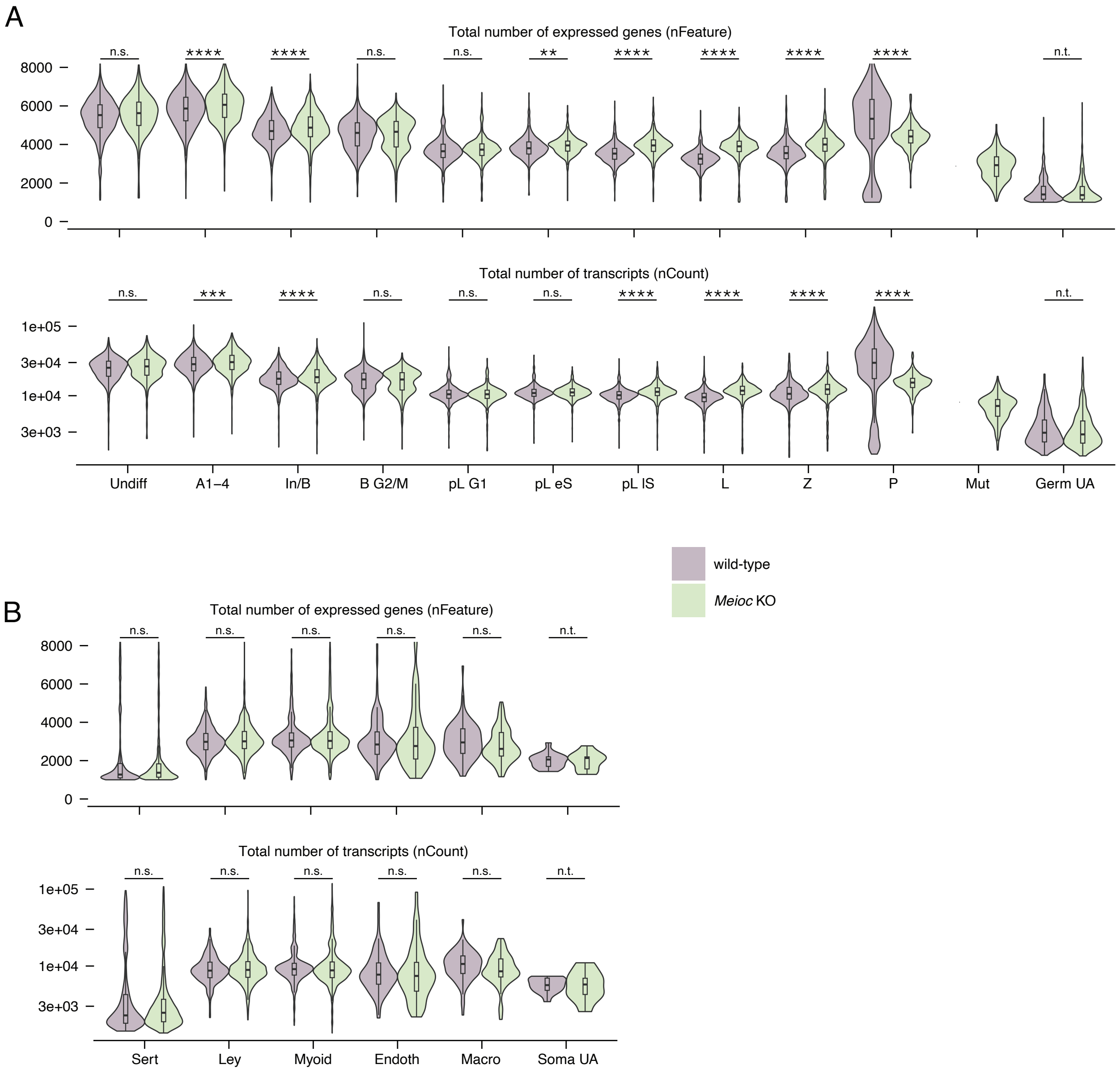
A
Total number of expressed genes (nFeature)
n.s.
****
****
n.s.
n.s.
**
****
****
****
****
n.t.
8000
6000
4000
2000
0
Total number of transcripts (nCount)
n.s.
***
****
n.s.
n.s.
n.s.
****
****
****
****
n.t.
1e+05
3e+04
1e+04
3e+03
Undiff
A1–4
In/B
B G2/M
pL G1
pL eS
pL lS
L
Z
P
Mut
Germ UA
wild-type
Meioc KO
B
Total number of expressed genes (nFeature)
n.s.
n.s.
n.s.
n.s.
n.s.
n.t.
8000
6000
4000
2000
0
Total number of transcripts (nCount)
n.s.
n.s.
n.s.
n.s.
n.s.
n.t.
1e+05
3e+04
1e+04
3e+03
Sert
Ley
Myoid
Endoth
Macro
Soma UA

**Figure S2: Features and counts in scRNA-seq data.**

**A:** Total number of genes expressed (nFeature) and transcripts detected (nCount) in wild-type and *Meioc* KO cells for germ cell clusters.
**B:** Total number of genes expressed (nFeature) and transcripts detected (nCount) in wild-type and *Meioc* KO cells for somatic clusters.

*, adj. $P<0.05$; **, adj. $P<0.01$; ***, adj. $P<0.001$; ****, adj. $P<0.0001$; n.s., not significant; n.t., not tested, as populations may represent sequencing artifacts.

### Figure S3



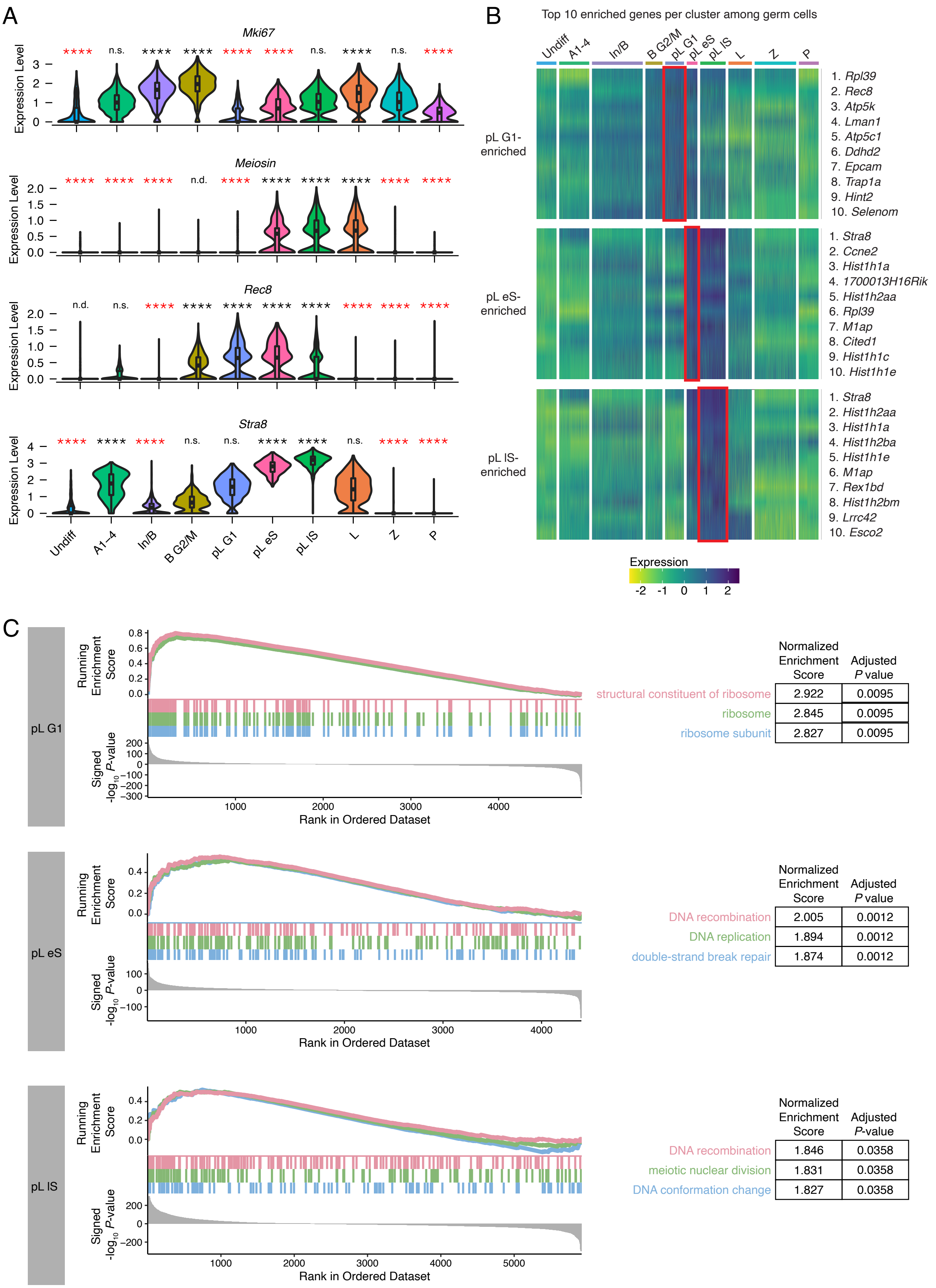


| | Normalized Enrichment Score | Adjusted P value |
|---|---|---|
| structural constituent of ribosome | 2.922 | 0.0095 |
| ribosome | 2.845 | 0.0095 |
| ribosome subunit | 2.827 | 0.0095 |

| | Normalized Enrichment Score | Adjusted P value |
|---|---|---|
| DNA recombination | 2.005 | 0.0012 |
| DNA replication | 1.894 | 0.0012 |
| double-strand break repair | 1.874 | 0.0012 |

| | Normalized Enrichment Score | Adjusted P-value |
|---|---|---|
| DNA recombination | 1.846 | 0.0358 |
| meiotic nuclear division | 1.831 | 0.0358 |
| DNA conformation change | 1.827 | 0.0358 |

**Figure S3: Distinguishing features among wild-type preleptotene clusters.**

**A:** Wild-type expression levels of *Mki67*, *Meiosin*, *Rec8*, and *Stra8* for all germ cell clusters. The pL G1 cluster exhibited reduced *Mki67*, whose fluctuations during the mitotic cell cycle reach a nadir in G1 phase. *Stra8* and *Meiosin*, which encode a heterodimer transcriptional activator required for the meiotic G1/S phase transition, are robustly expressed in the pL eS and lS clusters. *Rec8*, a meiosis-specific cohesin that is activated by retinoic acid independent of *Stra8* (Soh et al., 2015), is also enriched in pL clusters. Black and red asterisks designate enrichment or depletion, respectively, relative to all other germ cells. Clusters marked as "not done" (n.d.) did not meet expression thresholds set for statistical testing.
**B:** Heatmap of the top 10 enriched genes in the pL G1, eS, and lS clusters. The pL G1 cluster was enriched for *Rpl39* expression, consistent with observations that mitotic G1 phase is associated with high levels of ribosome biogenesis (Derenzini et al., 2005; Gómez-Herreros et al., 2013; Nosrati et al., 2014; Volarević et al., 2000). Replication-dependent histones were among the top 10 enriched genes in pL eS and lS clusters, confirming the S phase designation of these clusters.
**C:** Gene Set Enrichment Analysis (GSEA) for Gene Ontology (GO) terms for genes within each celltype cluster. The pL G1 cluster was enriched for ribosome-associated Gene Ontology terms, consistent with the high levels of ribosome biogenesis observed in mitotic G1 phase (Derenzini et al., 2005; Gómez-Herreros et al., 2013; Nosrati et al., 2014; Volarević et al., 2000). The preleptotene clusters in early and late S phase were enriched for expression of genes associated with GO term "DNA recombination," consistent with their S phase designations. Genes were ranked by the signed $-\log_{10}(P$ value) for their level of enrichment/depletion within each cluster relative to all other germ cell clusters. Top three enriched GO terms for each cluster shown.

*, adj. $P<0.05$; **, adj. $P<0.01$; ***, adj. $P<0.001$; ****, adj. $P<0.0001$; n.s., not significant; n.d., not detected (transcript expressed in <25% cells in each population being compared).

Figure S4



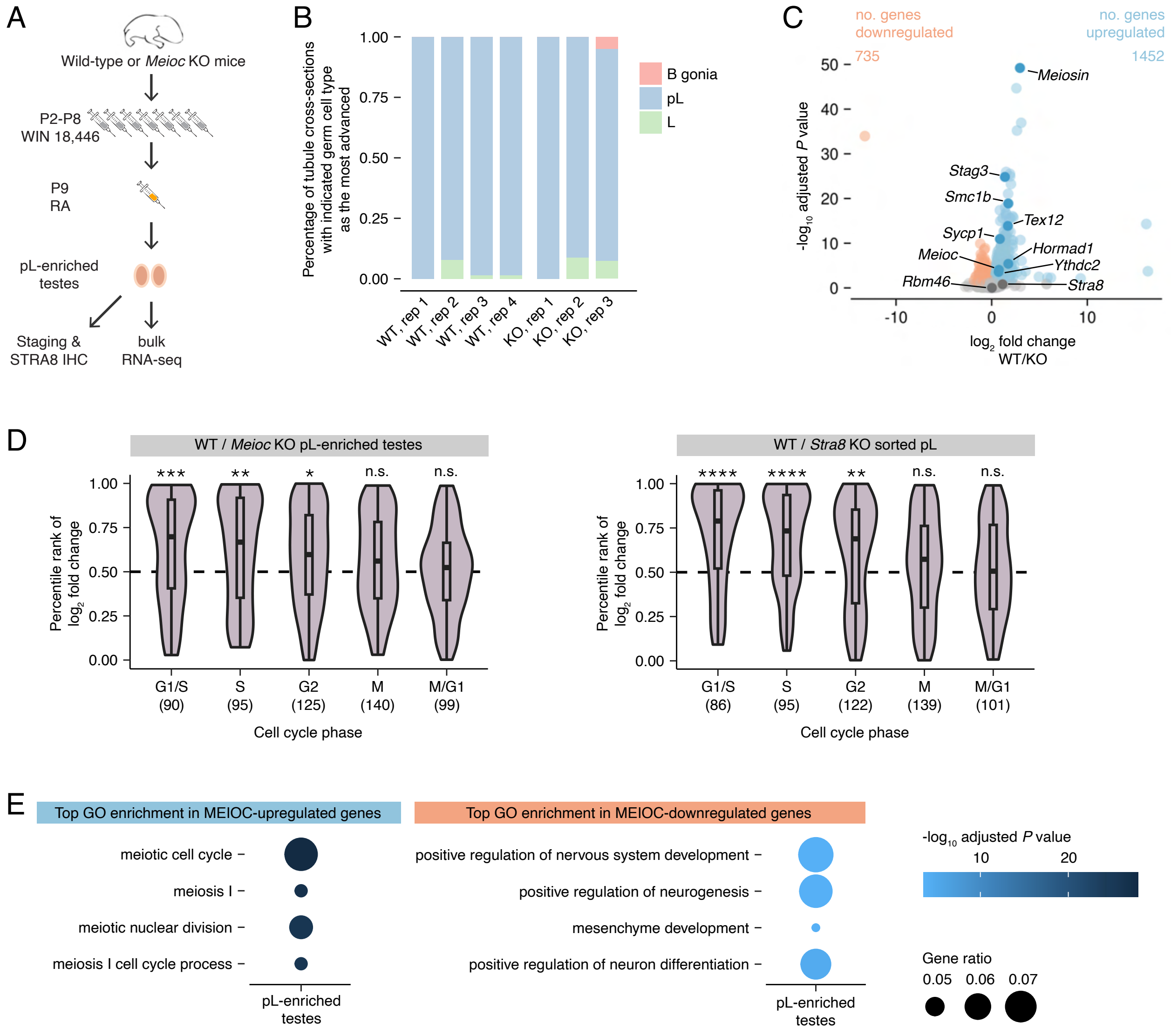
A
Wild-type or Meioc KO mice
P2-P8
WIN 18,446
P9
RA
pL-enriched
testes
Staging &
STRA8 IHC
bulk
RNA-seq
B
Percentage of tubule cross-sections with indicated germ cell type as the most advanced
1.00
0.75
0.50
0.25
0.00
WT, rep 1
WT, rep 2
WT, rep 3
WT, rep 4
KO, rep 1
KO, rep 2
KO, rep 3
B gonia
pL
L
C
no. genes downregulated
735
no. genes upregulated
1452
-log10 adjusted P value
log2 fold change WT/KO
Meiosin
Stag3
Smc1b
Tex12
Sycp1
Hormad1
Meioc
Ythdc2
Rbm46
Stra8
D
WT / Meioc KO pL-enriched testes
Percentile rank of log2 fold change
***
**
*
n.s.
n.s.
G1/S (90)
S (95)
G2 (125)
M (140)
M/G1 (99)
Cell cycle phase
WT / Stra8 KO sorted pL
****
****
**
n.s.
n.s.
G1/S (86)
S (95)
G2 (122)
M (139)
M/G1 (101)
Cell cycle phase
E
Top GO enrichment in MEIOC-upregulated genes
meiotic cell cycle
meiosis I
meiotic nuclear division
meiosis I cell cycle process
pL-enriched testes
Top GO enrichment in MEIOC-downregulated genes
positive regulation of nervous system development
positive regulation of neurogenesis
mesenchyme development
positive regulation of neuron differentiation
pL-enriched testes
-log10 adjusted P value
10
20
Gene ratio
0.05 0.06 0.07

**Figure S4: MEIOC-dependent program defined by bulk RNA-seq analysis of preleptotene-enriched testes.**

**A:** Developmental synchronization of spermatogenesis for bulk RNA-seq analysis. IHC, immunohistochemistry; RA, retinoic acid.
**B:** Enrichment for preleptotene spermatocytes in developmentally synchronized testes.
**C:** Differential expression of bulk RNA-seq data. Blue marks genes that are upregulated in response to MEIOC ($\log_2$ fold change>0 and adj. *P*<0.05). Orange marks genes that are downregulated in response to MEIOC ($\log_2$ fold change<0 and adj. *P*<0.05). Rbm46 and Stra8 are highlighted in dark gray; *Meioc*, *Ythdc2*, and *Meiosin* expression are highlighted in dark blue. Other dark blue genes represent those that fall under meiosis-associated GO terms from panel E.
**D:** Cell cycle analysis of differential expression from bulk RNA-seq data. MEIOC increases abundance of genes associated with G1/S, S, and G2 phases, similar to STRA8. WT and *Stra8* KO bulk RNA-seq data from sorted preleptotenes were reanalyzed from Kojima et al., 2019.
**E:** Top four GO gene lists enriched in MEIOC-upregulated and -downregulated genes.

*, adj. *P*<0.05; **, adj. *P*<0.01; ***, adj. *P*<0.001; adj. *P*<0.0001; n.s., not significant.

Figure S5



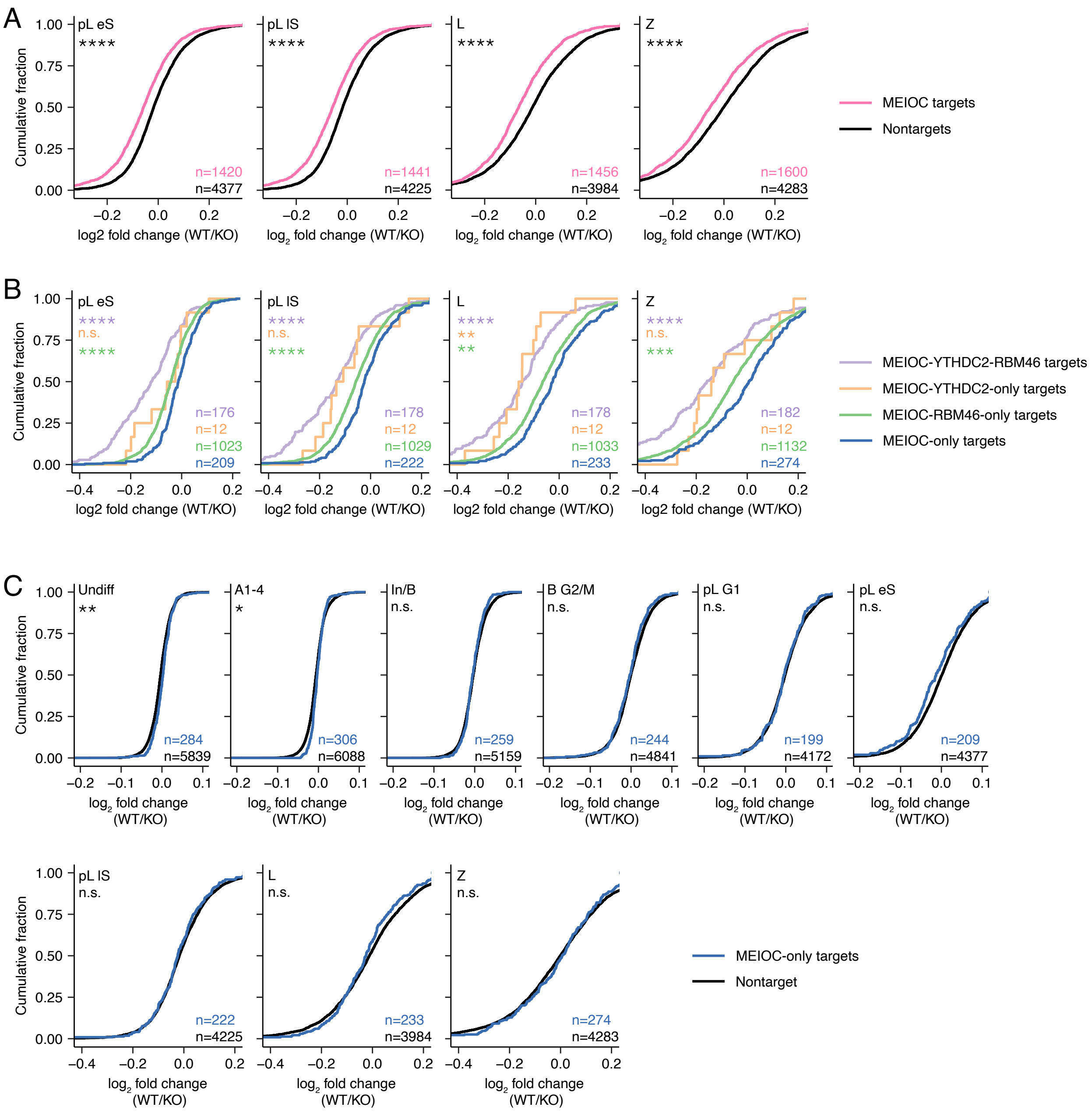
A
pL eS
pL lS
L
Z
****
Cumulative fraction
log2 fold change (WT/KO)
n=1420
n=4377
n=1441
n=4225
n=1456
n=3984
n=1600
n=4283
MEIOC targets
Nontargets
B
n.s.
**
***
n=176
n=12
n=1023
n=209
n=178
n=12
n=1029
n=222
n=178
n=12
n=1033
n=233
n=182
n=12
n=1132
n=274
MEIOC-YTHDC2-RBM46 targets
MEIOC-YTHDC2-only targets
MEIOC-RBM46-only targets
MEIOC-only targets
C
Undiff
A1-4
In/B
B G2/M
pL G1
n=284
n=5839
n=306
n=6088
n=259
n=5159
n=244
n=4841
n=199
n=4172
n=209
n=4377
n=222
n=4225
n=233
n=3984
n=274
n=4283
MEIOC-only targets
Nontarget

**Figure S5: MEIOC destabilizes its targets, based on scRNA-seq analysis.**

**A:** Cumulative fraction of $\log_2$ fold changes in transcript abundance in response to MEIOC (WT/ *Meioc* KO), in MEIOC targets compared to nontargets for pL eS, pL lS, L, and Z clusters. Plots for other clusters shown in Figure 2B.
**B:** Cumulative fraction of $\log_2$ fold changes in transcript abundance in response to MEIOC (WT/ *Meioc* KO), in MEIOC-YTHDC2-RBM46 targets, MEIOC-RBM46-only targets, MEOIC-YTHDC2-only targets, and MEIOC-only targets for pL eS, pL lS, L, and Z clusters. Asterisks represent comparison of color-matched target set to MEIOC-only targets. Plots for other clusters shown in Figure 2E.
**C:** Cumulative fraction of $\log_2$ fold changes in transcript abundance in response to MEIOC (WT/ *Meioc* KO), in MEIOC-only targets compared to nontargets for all germ cell clusters.

*, adj. $P$<0.05; **, adj. $P$<0.01; ***, adj. $P$<0.001; ****, adj. $P$<0.0001; n.s., not significant.

Figure S6



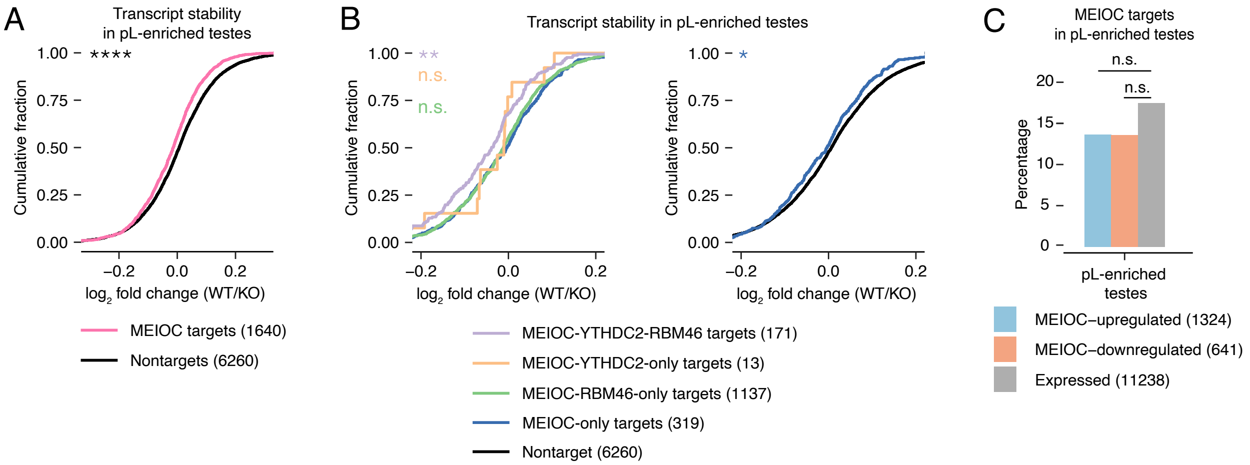

A
Transcript stability in pL-enriched testes
****
Cumulative fraction
1.00
0.75
0.50
0.25
0.00
−0.2
0.0
0.2
log2 fold change (WT/KO)
MEIOC targets (1640)
Nontargets (6260)
B
Transcript stability in pL-enriched testes
**
n.s.
n.s.
*
MEIOC-YTHDC2-RBM46 targets (171)
MEIOC-YTHDC2-only targets (13)
MEIOC-RBM46-only targets (1137)
MEIOC-only targets (319)
Nontarget (6260)
C
MEIOC targets in pL-enriched testes
Percentage
20
15
10
5
0
pL-enriched testes
MEIOC–upregulated (1324)
MEIOC–downregulated (641)
Expressed (11238)

**Figure S6: MEIOC destabilizes its targets, based on bulk RNA-seq analysis of preleptotene-enriched testes.**

**A:** Cumulative distribution of log2 fold change (WT/ *Meioc* KO) in transcript stability for MEIOC targets compared to nontargets. Transcript stability was estimated from bulk RNA-seq data of preleptotene-enriched testes.
**B:** Cumulative distribution of $_{\log 2}$ fold change (WT/ *Meioc* KO) in transcript stability for MEIOC-YTHDC2-RBM46 targets, MEIOC-RBM46-only targets, MEIOC-YTHDC2-only targets, and MEIOC-only targets (left), with asterisks representing comparison of color-matched target set to MEIOC-only targets. Cumulative distribution of $\log_2$ fold change (WT/ *Meioc* KO) in transcript stability for MEIOC-only targets compared to nontargets (right), with asterisk representing comparison between two groups.
**C:** Percentage of MEIOC targets among MEIOC-upregulated, MEIOC-downregulated, and expressed genes, defined by bulk RNA-seq analysis of preleptotene-enriched testes.

*, adj. $P$<0.05; **, adj. $P$<0.01; ***, adj. $P$<0.001; ****, adj. $P$<0.0001; n.s., not significant.

Figure S7



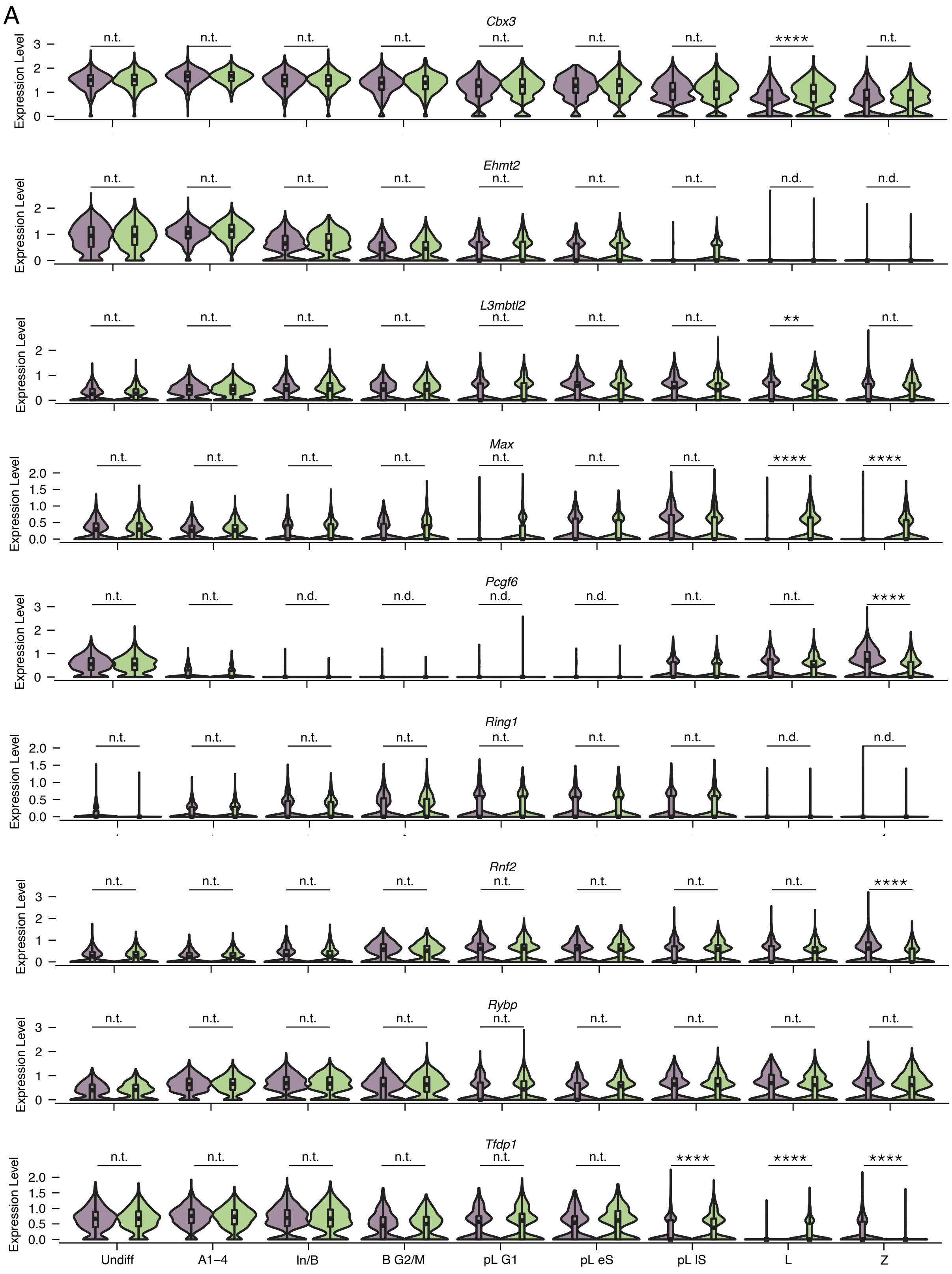
A
Cbx3
Ehmt2
L3mbtl2
Max
Pcgf6
Ring1
Rnf2
Rybp
Tfdp1
Expression Level
n.t.
n.d.
****
**
Undiff
A1–4
In/B
B G2/M
pL G1
pL eS
pL lS
L
Z

**Figure S7: Expression of PRC1.6 subunits in wild-type and *Meioc*-null cells from scRNA-seq data of P15 testes.**

**A:** Expression levels of transcripts for PRC1.6 subunits. Clusters marked as “not done” (n.d.) did not meet expression thresholds set for statistical testing.

*, adj. P<0.05; **, adj. *P*<0.01; ***, adj. *P*<0.001; ****, adj. *P*<0.0001; n.s., not significant; n.t., not tested (comparison was excluded from statistical testing because $\log_2$ fold change> -0.1 and <0.1); n.d., not detected (transcript expressed in <25% cells in each population being compared).

Figure S8



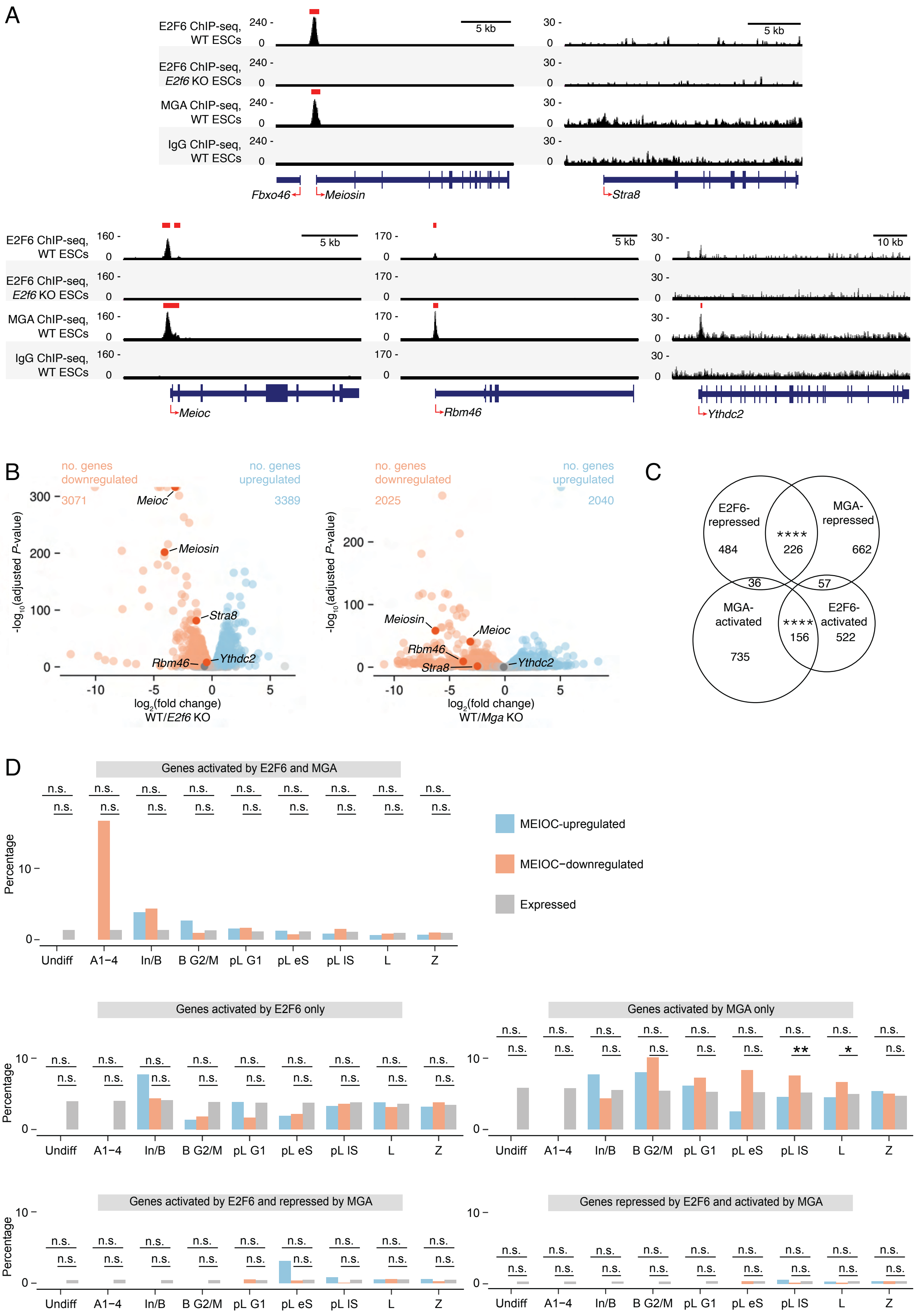
A
E2F6 ChIP-seq, WT ESCs
E2F6 ChIP-seq, E2f6 KO ESCs
MGA ChIP-seq, WT ESCs
IgG ChIP-seq, WT ESCs
5 kb
10 kb
Fbxo46
Meiosin
Stra8
Meioc
Rbm46
Ythdc2
B
no. genes downregulated
no. genes upregulated
3071
3389
2025
2040
-log10(adjusted P-value)
log2(fold change) WT/E2f6 KO
log2(fold change) WT/Mga KO
C
E2F6-repressed
MGA-repressed
MGA-activated
E2F6-activated
484
226
662
36
57
735
156
522
D
Genes activated by E2F6 and MGA
Genes activated by E2F6 only
Genes activated by MGA only
Genes activated by E2F6 and repressed by MGA
Genes repressed by E2F6 and activated by MGA
Percentage
MEIOC-upregulated
MEIOC−downregulated
Expressed
Undiff
A1−4
In/B
B G2/M
pL G1
pL eS
pL lS
L
Z
n.s.

**Figure S8: MEIOC-YTHDC2-RBM46's repression of *E2f6* and *Mga* mRNA relieves E2F6- and MGA-mediated transcriptional repression.**

**A:** Input-subtracted E2F6 ChIP-seq signal from wild-type and E2F6 KO ESCs as well as normalized MGA and IgG ChIP-seq signal from wild-type ESCs at the promoters of *Meiosin*, *Stra8*, *Meioc*, *Rbm46*, and *Ythdc2*. Called peaks are marked by a red bar above. Transcriptional start sites are marked by red arrows. E2F6 ChIP-seq data were reanalyzed from Dahlet et al., 2021; MGA ChIP-seq data were reanalyzed from Stielow et al., 2018.
**B:** E2F6-dependent and MGA-dependent differential expression program in mouse embryonic stem cells. RNA-seq data were reanalyzed from Dahlet et al., 2021 and Qin et al., 2021. $\mathrm{Log}_2$ fold change was defined as WT/KO. Genes upregulated or downregulated by E2F6 or MGA are shown in blue and orange, respectively. Gray represents genes that do not change expression in response to E2F6 or MGA.
**C:** Overlap of E2F6-repressed and -activated genes with MGA-repressed and -activated genes, identified from reanalysis of published ChIP-seq and RNA-seq datasets from mouse embryonic stem cells.
**D:** Percentage of E2F6- and/or MGA-regulated genes among MEIOC-upregulated, MEIOC-downregulated, and expressed genes, as identified via scRNA-seq analysis. Each E2F6/MGA gene set was tested for enrichment among MEIOC-upregulated genes and MEIOC-downregulated genes relative to expressed genes. E2F6-regulated genes and MGA-regulated genes were identified as shown in panel C.

*, adj. $P<0.05$; **, adj. $P<0.01$; ***, adj. $P<0.001$; ****, adj. $P<0.0001$; n.s., not significant.

### Figure S9



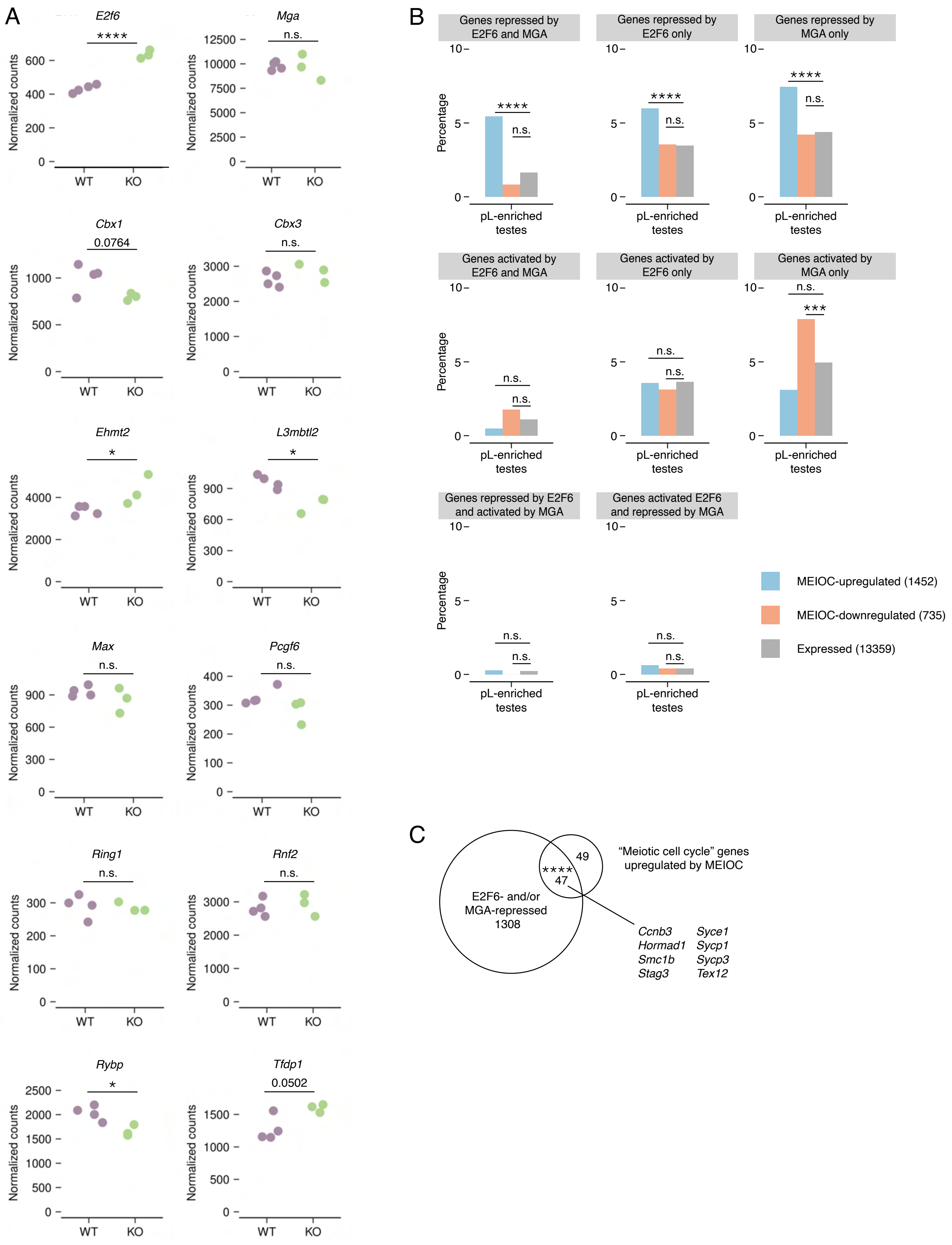
A
E2f6
****
Normalized counts
WT
KO
Mga
n.s.
Cbx1
0.0764
Cbx3
n.s.
Ehmt2
*
L3mbtl2
*
Max
n.s.
Pcgf6
n.s.
Ring1
n.s.
Rnf2
n.s.
Rybp
*
Tfdp1
0.0502
B
Genes repressed by E2F6 and MGA
Genes repressed by E2F6 only
Genes repressed by MGA only
Genes activated by E2F6 and MGA
Genes activated by E2F6 only
Genes activated by MGA only
Genes repressed by E2F6 and activated by MGA
Genes activated E2F6 and repressed by MGA
Percentage
pL-enriched testes
****
***
n.s.
MEIOC-upregulated (1452)
MEIOC-downregulated (735)
Expressed (13359)
C
E2F6- and/or MGA-repressed 1308
****
47
49
"Meiotic cell cycle" genes upregulated by MEIOC
Ccnb3
Hormad1
Smc1b
Stag3
Syce1
Sycp1
Sycp3
Tex12

**Figure S9: MEIOC-YTHDC2-RBM46 repression of *E2f6* and *Mga* mRNA relieves E2F6- and MGA-mediated transcriptional repression, based on bulk RNA-seq analysis.**

**A:** Normalized counts in WT vs. *Meioc* KO bulk RNA-seq analysis of preleptotene-enriched testes for *E2f6*, *Mga*, and other subunits of PRC1.6.
**B:** Percentage of E2F6-repressed and -activated, as well as MGA-repressed and -activated, genes among MEIOC-upregulated, MEIOC-downregulated, and expressed genes from bulk RNA-seq analysis of preleptotene-enriched testes. E2F6 and MGA targets as defined in Figure S8C.
**C:** Overlap between E2F6- and/or MGA-repressed genes and the “meiotic cell cycle” genes upregulated by MEIOC in preleptotene-enriched testes.

*, adj. $P<0.05$; **, adj. $P<0.01$; ***, adj. $P<0.001$; ****, adj. $P<0.0001$; n.s., not significant.

Figure S10



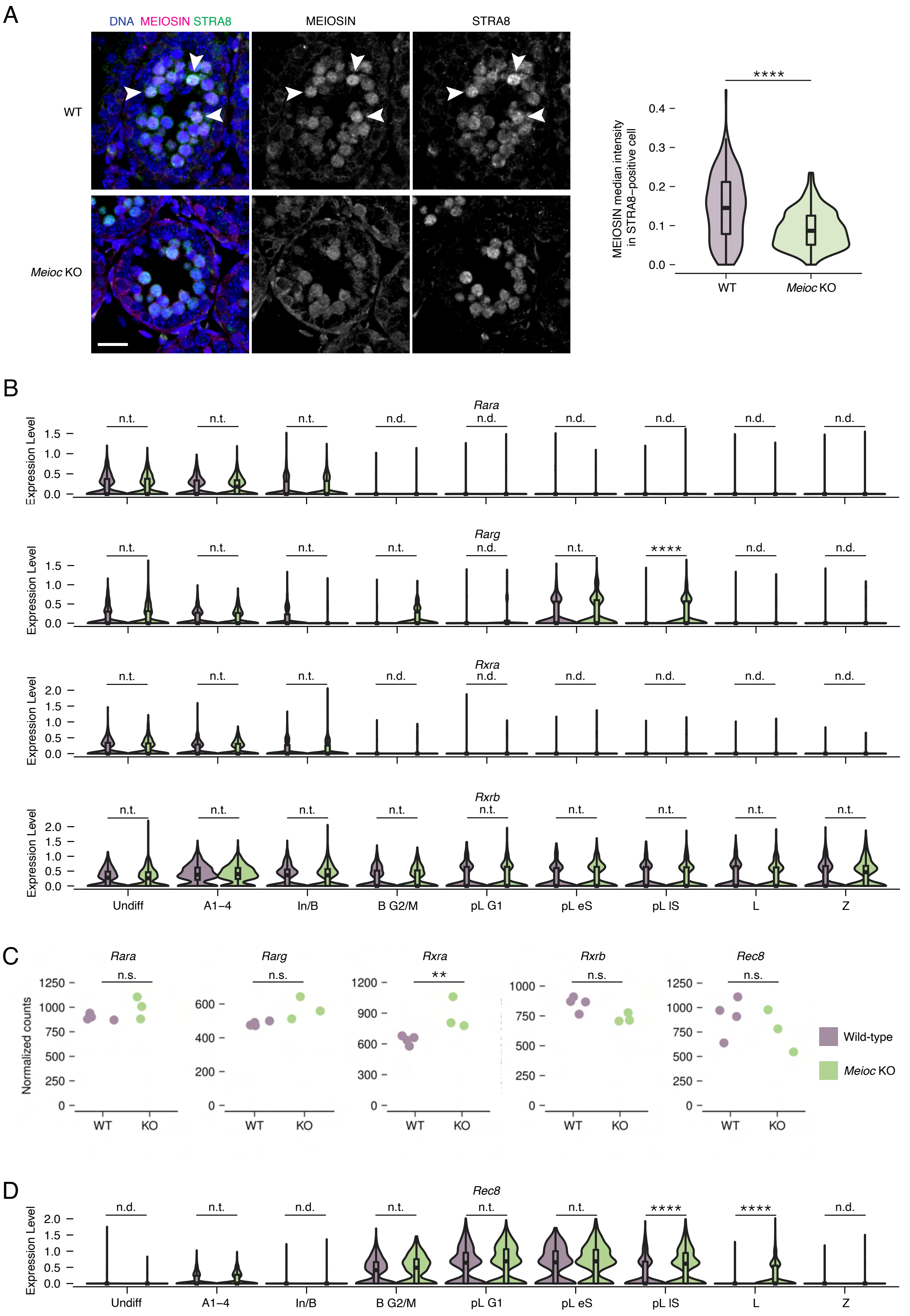
A
DNA MEIOSIN STRA8
MEIOSIN
STRA8
WT
Meioc KO
MEIOSIN median intensity in STRA8–positive cell
****
B
Rara
Rarg
Rxra
Rxrb
Expression Level
Undiff
A1–4
In/B
B G2/M
pL G1
pL eS
pL lS
L
Z
C
Rara
Rarg
Rxra
Rxrb
Rec8
Normalized counts
WT
KO
Wild-type
Meioc KO
D
Rec8

**Figure S10: MEIOC increases Meiosin gene expression in response to retinoic acid.**

**A:** MEIOSIN protein expression in STRA8-positive preleptotene spermatocytes from wild-type and *Meioc* KO P10 testes. Arrowheads represent STRA8-positive spermatocytes with particularly robust MEIOSIN that are absent from Meioc KO testes. Graph represents MEIOSIN median intensity from three WT and *Meioc* KO littermate pairs. 75 cells were quantified per testis. Scale bar = 20 um.
**B:** Expression levels of *Rar* and *Rxr* genes in wildtype vs. *Meioc*-null cells in all germ cell clusters identified in the scRNA-seq analysis. *Rarb* and *Rxrg* were not detected as expressed. Color legend same as in panel C. Clusters marked as “not done” (n.d.) did not meet expression thresholds set for statistical testing.
**C:** Normalized counts of *Rar* and *Rxr* genes as well as *Rec8* in bulk RNA-seq analysis of wildtype vs. *Meioc*-null preleptotene-enriched testes. *Rarb* and *Rxrg* were not detected as expressed.
**D:** Expression levels of *Rec8* in wildtype vs. *Meioc*-null cells in all germ cell clusters identified in the scRNA-seq analysis. Color legend same as in panel C. Clusters marked as “not done” (n.d.) did not meet expression thresholds set for statistical testing.

*, adj. $P$<0.05; **, adj. $P$<0.01; ***, adj. $P$<0.001; ****, adj. $P$<0.0001; n.s., not significant; n.t., not tested (comparison was excluded from statistical testing because $\log_2$ fold change> -0.1 and <0.1); n.d., not detected (transcript expressed in <25% cells in each population being compared).

Figure S11



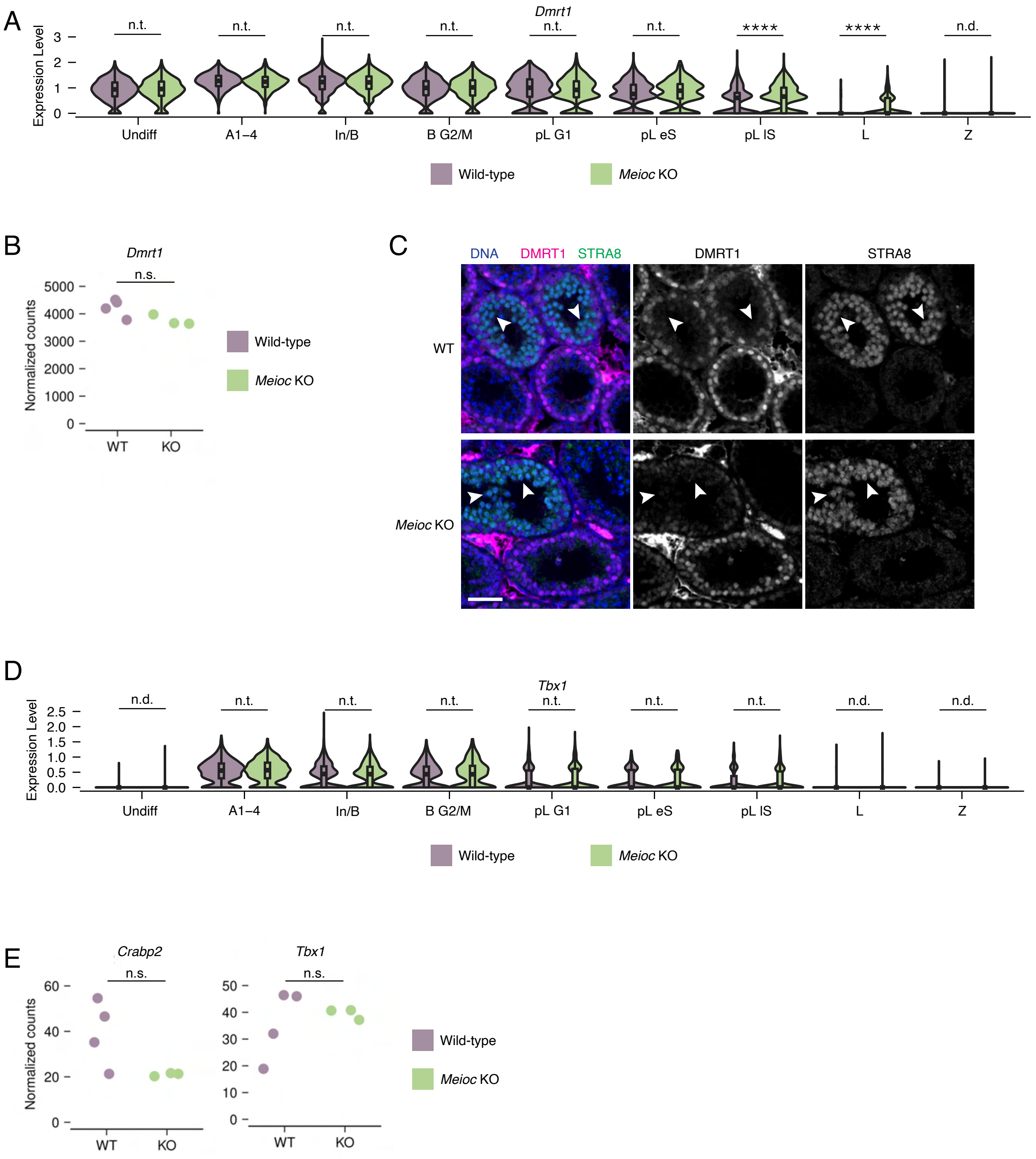
A
Dmrt1
Expression Level
n.t.
n.t.
n.t.
n.t.
n.t.
n.t.
****
****
n.d.
Undiff
A1–4
In/B
B G2/M
pL G1
pL eS
pL IS
L
Z
Wild-type
Meioc KO
B
Dmrt1
n.s.
Normalized counts
WT
KO
Wild-type
Meioc KO
C
DNA DMRT1 STRA8
DMRT1
STRA8
WT
Meioc KO
D
Tbx1
Expression Level
n.d.
n.t.
n.t.
n.t.
n.t.
n.t.
n.t.
n.d.
n.d.
Undiff
A1–4
In/B
B G2/M
pL G1
pL eS
pL IS
L
Z
Wild-type
Meioc KO
E
Crabp2
Tbx1
n.s.
n.s.
Normalized counts
WT
KO
WT
KO
Wild-type
Meioc KO

**Figure S11: MEIOC does not affect DMRT1 expression or DMRT1-dependent signaling during the mitosis-to-meiosis transition.**

**A:** Expression levels of *Dmrt1* in wildtype vs. *Meioc*-null cells in all germ cell clusters identified in the scRNA-seq analysis. Cluster marked as "not done" (n.d.) did not meet expression thresholds set for statistical testing.
**B:** Normalized counts of *Dmrt1* in bulk RNA-seq analysis of wildtype vs. *Meioc*-null preleptotene-enriched testes.
**C:** DMRT1 protein was not detected in STRA8-positive preleptotene spermatocytes (white arrowheads) in both wild-type and *Meioc* KO P15 testes. Scale bar = 40 um.
**D:** Expression levels of DMRT1-regulated gene *Tbx1* in wildtype vs. *Meioc*-null cells in all germ cell clusters identified in the scRNA-seq analysis. Clusters marked as "not done" (n.d.) did not meet expression thresholds set for statistical testing. *Crabp2* was not detected as expressed.
**E:** Normalized counts of DMRT1-regulated genes *Crabp2* and *Tbx1* in bulk RNA-seq analysis of wildtype vs. *Meioc*-null preleptotene-enriched testes.

*, adj. $P<0.05$; **, adj. $P<0.01$; ***, adj. $P<0.001$; ****, adj. $P<0.0001$; n.s., not significant; n.t., not tested (comparison was excluded from statistical testing because $\log_2$ fold change> -0.1 and <0.1); n.d., not detected (transcript expressed in <25% cells in each population being compared).

Figure S12



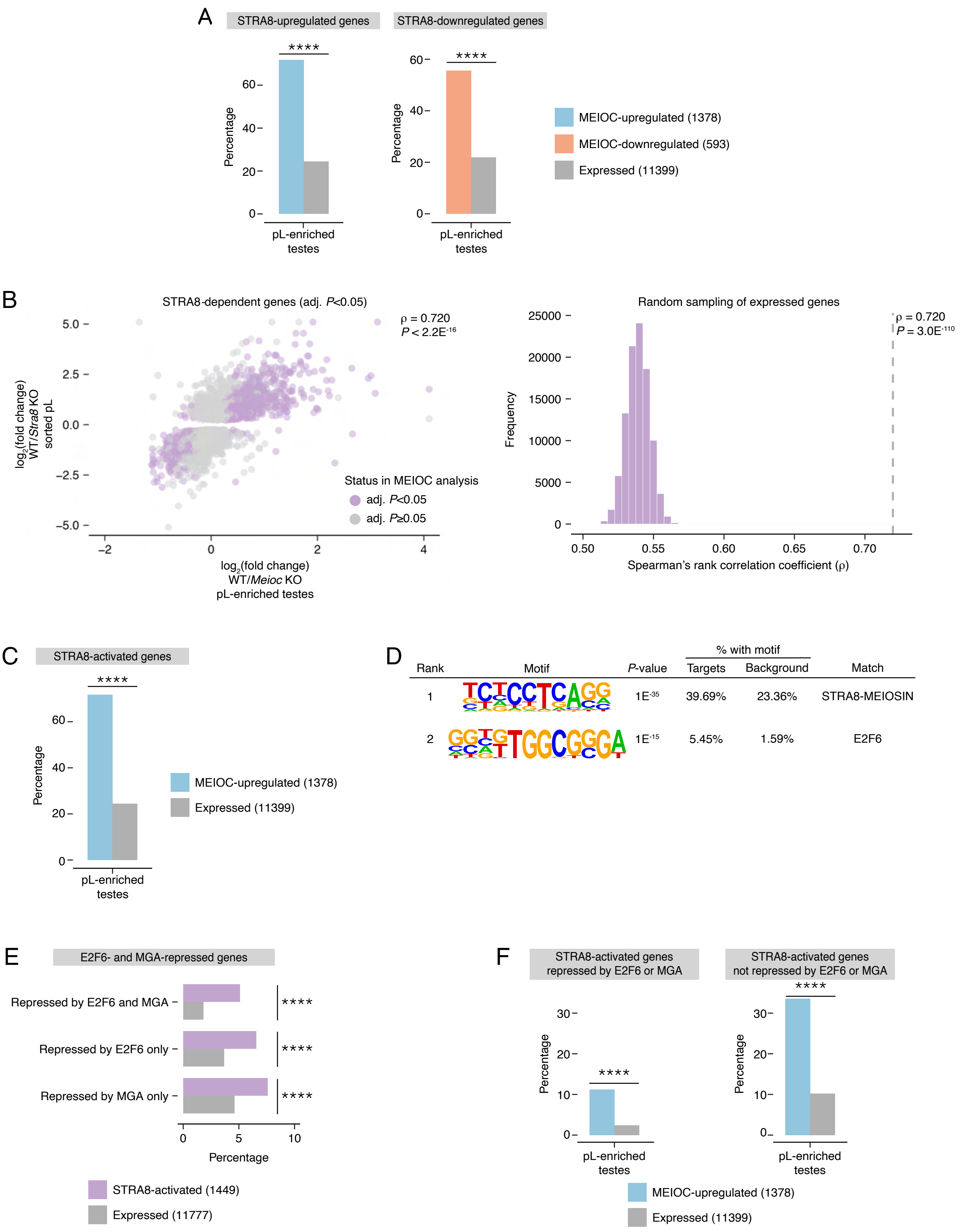
A
STRA8-upregulated genes
STRA8-downregulated genes
****
Percentage
pL-enriched testes
MEIOC-upregulated (1378)
MEIOC-downregulated (593)
Expressed (11399)
B
STRA8-dependent genes (adj. P<0.05)
ρ = 0.720
P < 2.2E-16
log2(fold change) WT/Stra8 KO sorted pL
log2(fold change) WT/Meioc KO pL-enriched testes
Status in MEIOC analysis
adj. P<0.05
adj. P≥0.05
Random sampling of expressed genes
ρ = 0.720
P = 3.0E-110
Frequency
Spearman's rank correlation coefficient (ρ)
C
STRA8-activated genes
****
Percentage
pL-enriched testes
MEIOC-upregulated (1378)
Expressed (11399)
D
Rank
Motif
P-value
% with motif
Targets
Background
Match
1
1E-35
39.69%
23.36%
STRA8-MEIOSIN
2
1E-15
5.45%
1.59%
E2F6
E
E2F6- and MGA-repressed genes
Repressed by E2F6 and MGA
Repressed by E2F6 only
Repressed by MGA only
****
Percentage
STRA8-activated (1449)
Expressed (11777)
F
STRA8-activated genes repressed by E2F6 or MGA
STRA8-activated genes not repressed by E2F6 or MGA
****
Percentage
pL-enriched testes
MEIOC-upregulated (1378)
Expressed (11399)

**Figure S12: MEIOC's derepression of *Meiosin* gene expression activates the STRA8-MEIOSIN transcriptional program in bulk RNA-seq data of preleptotene-enriched testes.**

**A:** Percentage of STRA8-upregulated and -downregulated genes in MEIOC-upregulated, -downregulated, and expressed genes from bulk RNA-seq analysis of preleptotene-enriched testes. STRA8-upregulated and -downregulated genes were identified via reanalysis of bulk RNA-seq data from wild-type and *Stra8* KO sorted preleptotene spermatocytes from Kojima et al., 2019.
**B:** Left panel, correlation between MEIOC bulk RNA-seq analysis of preleptotene-enriched testes and STRA8 bulk RNA-seq analysis of sorted preleptotene spermatocytes. Analysis was limited to genes that were statistically dependent on STRA8 (adj. $P<0.05$). *P* value represents the probability that Spearman rho does not equal 0. Right panel, distribution of correlations for gene sets randomly sampled from genes expressed in the two bulk RNA-seq datasets. *P* value represents that probability of obtaining an equal or larger correlation by random sampling.
**C:** Percentage of STRA8-activated genes in MEIOC-upregulated and expressed genes from bulk RNA-seq analysis of preleptotene-enriched testes. STRA8-activated genes were identified as those genes with STRA8-bound promoters (as identified by Kojima et al. (2019) via STRA8-FLAG ChIP-seq in preleptotene-enriched testes) and upregulated by STRA8 (as identified by reanalysis of bulk RNA-seq data from wild-type and *Stra8* KO sorted preleptotene spermatocytes from Kojima et al., 2019).
**D.** Motif enrichment within promoters of MEIOC-upregulated genes from bulk RNA-seq analysis of preleptotene-enriched testes. The second ranked motif was matched to E2F6, in addition to other E2F proteins and SMAD3.
**E:** Percentage of genes repressed by both E2F6 and MGA; E2F6 only; and MGA only in STRA8-activated and all expressed genes. Genes repressed by E2F6 and MGA were defined in mouse embryonic stem cells as shown in Figure S8C.
**F:** Percentage of genes that are activated by STRA8 and repressed by E2F6 or MGA; and activated by STRA8 but not repressed by E2F6 or MGA among MEIOC-upregulated and expressed genes from bulk RNA-seq analysis of preleptotene-enriched testes. Genes repressed by E2F6 or MGA are defined as those that are repressed by both E2F6 and MGA; by E2F6 only; or by MGA only, as shown in Figure S8C.

*, adj. $P<0.05$; **, adj. $P<0.01$; ***, adj. $P<0.001$; ****, adj. $P<0.0001$; n.s., not significant.

### Figure S13



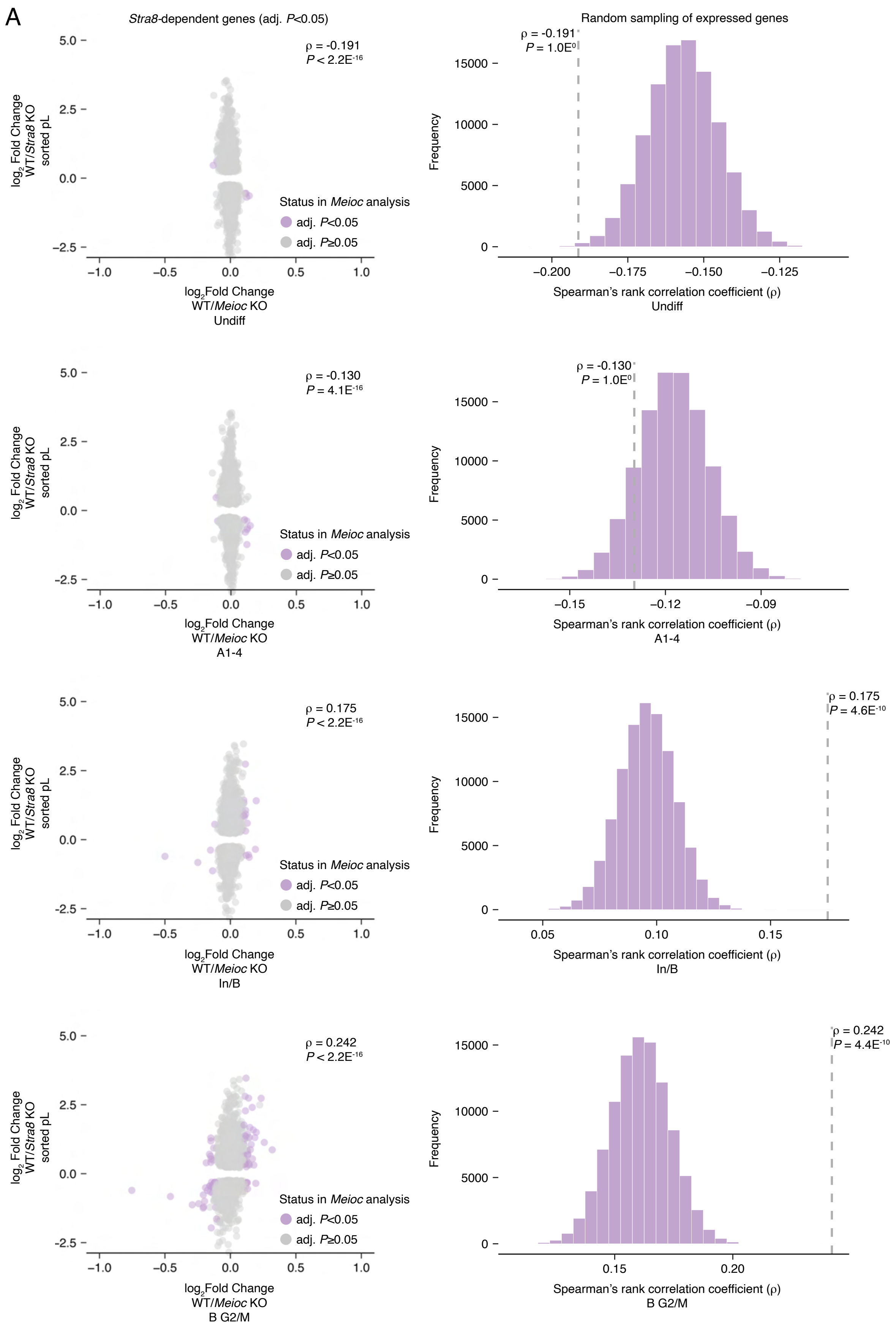
A
Stra8-dependent genes (adj. P<0.05)
Random sampling of expressed genes
ρ = -0.191
P < 2.2E-16
ρ = -0.191
P = 1.0E0
log2 Fold Change
WT/Stra8 KO
sorted pL
Frequency
Status in Meioc analysis
adj. P<0.05
adj. P≥0.05
log2Fold Change
WT/Meioc KO
Undiff
Spearman's rank correlation coefficient (ρ)
Undiff
ρ = -0.130
P = 4.1E-16
ρ = -0.130
P = 1.0E0
A1-4
A1-4
ρ = 0.175
P < 2.2E-16
ρ = 0.175
P = 4.6E-10
In/B
In/B
ρ = 0.242
P < 2.2E-16
ρ = 0.242
P = 4.4E-10
B G2/M
B G2/M

**Figure S13: Correlation between transcript abundance changes induced by MEIOC and STRA8: A1-4, In/B, and B G2/M clusters.**

**A:** Left panels, correlation between MEIOC scRNA-seq analysis of Undiff, A1-4, In/B, and B G2/M clusters and STRA8 bulk RNA-seq analysis of sorted preleptotene spermatocytes. Analysis was limited to genes that were statistically dependent on STRA8 (adj. P<0.05). *P* value represents the probability that Spearman rho does not equal 0. Right panels, distribution of correlations for gene sets randomly sampled from genes expressed in the scRNA-seq Undiff, A1-4, In/B, and B G2/M clusters and bulk RNA-seq sorted preleptotene spermatocytes. *P* value represents that probability of obtaining an equal or larger correlation by random sampling.

Figure S14



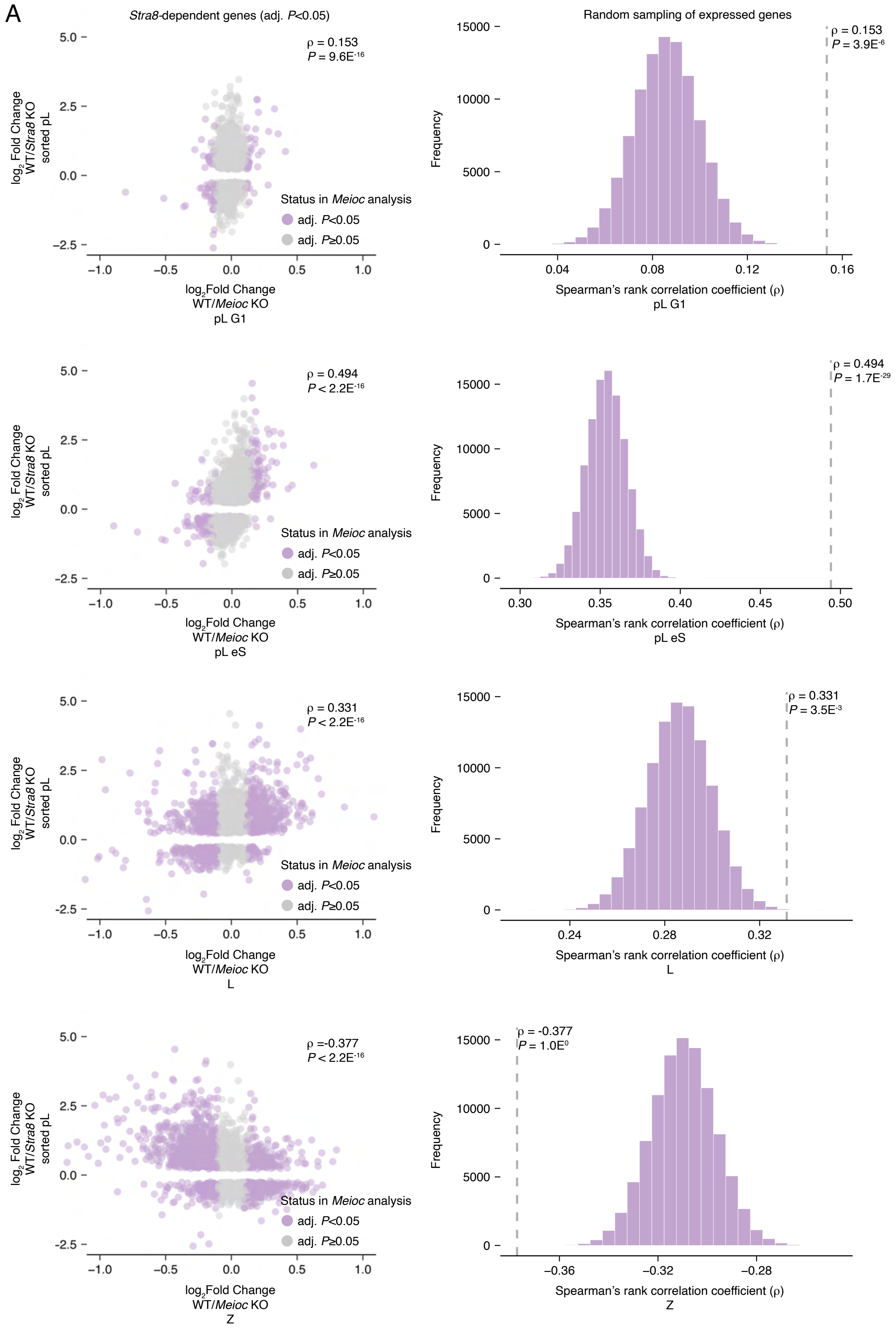

A
Stra8-dependent genes (adj. P<0.05)
Random sampling of expressed genes
ρ = 0.153
P = 9.6E-16
P = 3.9E-6
ρ = 0.494
P < 2.2E-16
P = 1.7E-29
ρ = 0.331
P = 3.5E-3
ρ =-0.377
ρ = -0.377
P = 1.0E0
log2 Fold Change WT/Stra8 KO sorted pL
log2Fold Change WT/Meioc KO
pL G1
pL eS
L
Z
Status in Meioc analysis
adj. P<0.05
adj. P≥0.05
Frequency
Spearman's rank correlation coefficient (ρ)

**Figure S14: Correlation between transcript abundance changes induced by MEIOC and STRA8: pL G1, pL eS, L, or Z clusters.**

A: Left panels, correlation between MEIOC scRNA-seq analysis of pL G1, pL eS, L, or Z clusters and STRA8 bulk RNA-seq analysis of sorted preleptotene spermatocytes. Analysis was limited to genes that were statistically dependent on STRA8 (adj. $P<0.05$). *P* value represents the probability that Spearman rho does not equal 0. Right panels, distribution of correlations for gene sets randomly sampled from genes expressed in the scRNA-seq pL G1, pL eS, L, or Z clusters and bulk RNA-seq sorted preleptotene spermatocytes. *P* value represents that probability of obtaining an equal or larger correlation by random sampling.

**Table S1: Supplementary table and stats by figure panel**

**Table S2: scRNA-seq analysis of WT germ cell clusters from P15 testis**

**Table S3: scRNA-seq differential expression analysis (DEA)**

**Table S4: scRNA-seq DEA WT v. KO Gene Ontology analysis**

**Table S5: Associated cell cycle phase of MEIOC-downregulated genes**

**Table S6: MEIOC, YTHDC2, and RBM46 target analysis**

**Table S7: mRNA stability and transcriptional rate from bulk RNA-seq data**

**Table S8: Number of cells per scRNA-seq cluster in P15 testes**

**Table S9: bulk RNA-seq differential expression analysis, WT vs. *Meioc* KO**

**Table S10: Cell cycle analysis of bulk RNA-seq *Meioc* and *Stra8* datasets**

**Table S11: bulk RNA-seq Gene Ontology analysis**

**Table S12: Reanalysis of E2f6 and Mga data from embryonic stem cells**

**Table S13: MEIOSIN quantification in WT v Meioc KO**
